# Identification of allo- or orthosteric VHH/single-domain antibodies that enhance or block pathogen binding to Siglec-1 on dendritic cells

**DOI:** 10.64898/2025.12.19.695420

**Authors:** H.J. Brink, A.J. Affandi, S. Man, N.C.C.T. Dekker, E.A. Große Wichtrup, R.G. Bouma, N. Seyed Toutounchi, V.A.L. Konijn, J.G.C. Stolwijk, J.L. van Hamme, K. Olesek, D.A.M. Heijnen, A. Fish, A. Moutsiopoulou, P.H.N. Celie, G. Dekkers, R. Heukers, E. Mastrobattista, M.J. Smit, Y. van Kooyk, A.P. Heikema, N.A. Kootstra, T.B.H. Geijtenbeek, J.M.M. den Haan

## Abstract

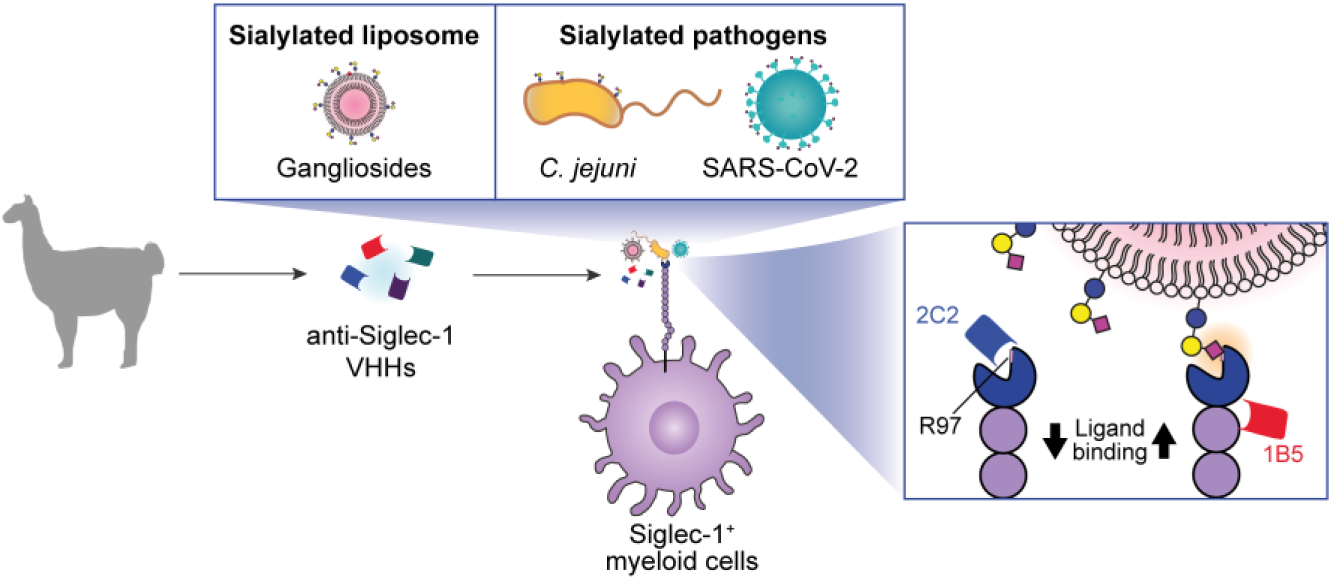

**Background:** Siglec-1 (Sialoadhesin/CD169) is expressed on myeloid cells and plays a key role in host defences by capturing incoming sialylated-pathogens such as *Campylobacter jejuni*. However, binding to Siglec-1 has also been exploited by pathogens such as SARS-CoV-2 for further dissemination.

**Results:** Here we identified high-affinity VHHs also known as single-domain antibodies or Nanobodies that bind to Siglec-1 and allo- or orthosterically modulate ligand binding. VHH 2C2 was shown to bind directly to the ligand binding site of Siglec-1 and blocked binding of ganglioside liposomes and *Campylobacter jejuni* to monocyte-derived dendritic cells (moDCs) and *ex vivo* Siglec-1^+^ DCs. VHH 2C2 also blocked SARS-CoV-2 binding of moDCs. In contrast, the VHHs 1B5 and 1C1 interacted with Siglec-1 outside the ligand binding site and acted as positive allosteric modulators of Siglec-1 ligand interactions, as was illustrated by increased ganglioside liposome and *Campylobacter jejuni* binding by moDCs. Our data suggests that mechanistically, the VHH 1B5 and 1C1 interfere with the *cis*-binding sialic acids present on the Siglec-1-expressing cell and thereby enhance *trans*-interactions with ligands.

**Conclusion:** In conclusion, we have isolated VHH that enhance or block Siglec-1 ligand binding to a variety of sialylated-pathogens enabling further interrogation of Siglec-1 function. Moreover, unlike conventional blocking antibodies targeting specific pathogens, Siglec-1 binding VHH could potentially serve as broad-spectrum pathogen blocking agents.

## Introduction

Sialic acid-binding immunoglobulin-like lectin 1 receptor (Siglec-1) also known as sialoadhesin or CD169, is expressed by macrophages and dendritic cells [1–8]. Siglec-1 is a large (182 kDa) type 1 membrane receptor that structurally consists of a N-terminal V-set immunoglobulin ligand binding domain, 16 C2-set immunoglobulin domains and a transmembrane domain with a short cytoplasmic tail that lacks an immunoreceptor tyrosine-based inhibitory motif (ITIM) typically found in other Siglecs [9, 10]. Siglec-1 binds to sialic acid N-acetylneuraminic acid (Neu5Ac) with a preference for α(2, 3)-linkage to O- and N-glycans on glycoproteins and glycolipids such as gangliosides. The affinity of ligands binding to Siglec-1 is low, with K_d_ values in the mM range, however binding of multivalent low-affinity ligands creates high-avidity interactions [11].

Siglec-1 mediates adhesion to sialic acid-expressing pathogens as well as host cells such as innate lymphocytes, neutrophils, DCs and regulatory T-cells [9, 12–14]. Due to its large size Siglec-1 extends far away from the cell membrane, enabling *trans-*interactions with sialic acids expressed on other cells while also capable of engaging ligands in *cis*-interactions [15–18]. Moreover, the extended size allows the capture of circulating pathogens in the lymph node subcapsular sinus and splenic marginal sinus which prevents further dissemination [19–22].

Known pathogens captured by Siglec-1 range from enveloped viruses including HIV-1, Ebola, and SARS-CoV-2 to bacteria such as *Neisseria meningitidis* and *Campylobacter jejuni* (*C. jejuni*) [5, 23–30]. In the case of HIV-1 and Ebola, Siglec-1 binds to GM1a and GM3 gangliosides that are incorporated into the viral membrane upon budding [24, 30]. HIV-1 and SARS-CoV-2 have developed strategies to exploit the capture by Siglec-1 to trans-infect neighbouring CD4^+^ T-cells or ACE2^+^ cells, enabling viral dissemination in the absence of a productive infection [24, 26, 31, 32]. Binding to Siglec-1 routes viruses to virus-containing-compartments that prevent degradation of virus and allow subsequent transmission to interacting cells [33, 34].

*C. jejuni* is a foodborne pathogen and one of the leading causes of bacterial gastroenteritis worldwide. A very small subset of patients develop Guillain-Barre Syndrome (GBS), due to the generation of antibodies against sialylated lipooligosaccharide (LOS) structures of *C. jejuni* that cross-react with host gangliosides such as GD1a, GM1 and GQ1b on peripheral nerves, resulting in paralysis [35–38]. GBS-associated *C. jejuni* strains preferentially bind to and are taken up by Siglec-1 macrophages and in a mouse model this Siglec-1-mediated uptake induces peripheral nerve neuropathy via the generation of ganglioside targeting auto-antibodies [39–42].

Considering the crucial role of Siglec-1 in capturing pathogens, our aim was to generate variable heavy domain of heavy chain-only antibodies (VHH) also known as single-domain antibodies or Nanobodies against human Siglec-1 that can modulate Siglec-1 ligand binding. VHH are small (∼15 kDa) and bind to their targets with high affinity [43]. VHH typically have long CDR3 loops that, combined with a relatively large paratope and small size, enable them to bind to epitopes inaccessible to conventional antibodies [43, 44]. VHH can act as conformational stabilizers, locking a target protein into a fixed state, enabling the interrogation of structure/function relationships [45, 46].

In this study, we generated high-affinity Siglec-1 targeting VHHs and characterized three clones (1B5, 1C1 and 2C2) for their capacity to bind and modulate Siglec-1 ligand interactions. VHH clones 1B5 and 1C1 bound Siglec-1 outside of the ligand binding site and have a predicted binding site on the interface between domain 1 and 2, whereas 2C2 bound directly to the ligand binding site of Siglec-1. We reveal that VHH 1B5 and 1C1 could positively modulate Siglec-1’s adhesive function, as observed by the enhancement of sialylated pathogen binding of *C. jejuni* and SARS-CoV-2 by cell lines and primary antigen presenting cells (APCs). In contrast, 2C2 blocked Siglec-1-dependent interactions with *C. jejuni* and SARS-CoV-2. We have, therefore, generated new tools that positively or negatively regulate Siglec-1 binding to various sialylated pathogens, which will help to further decipher Siglec-1’s function and could be explored as potential therapeutics.

## Material and Methods

### Cell Culture

BW-5147 cells overexpressing human Siglec-1 (BwSn), THP-1 cells overexpressing human Siglec-1 (TSn), parental THP-1 and BW-5147 (BW; [47]) cells were maintained in RPMI 1640 (Thermo Fisher Scientific) complete medium, containing 10% fetal calf serum (Biowest), 50 U/ml penicillin, 50 μg/ml streptomycin and 2 mM glutamine (all from Thermo Fisher Scientific) at 37˚C and 5% CO_2_.

### Generation of monocyte-Derived DCs (moDCs)

Monocytes were isolated using a Percoll gradient or CD14-magnetic beads (Miltenyi Biotec) and then were cultured for five to six days in RPMI 1640 complete medium (Thermo Fisher Scientific) containing 10% fetal calf serum (Biowest), 50 U/mL penicillin, 50 μg/mL streptomycin, and 2 mM glutamine (all from Thermo Fisher Scientific), in the presence of recombinant human IL-4 (500 U/mL) and GM-CSF (800 U/mL; both from Immunotools) at 37˚C and 5% CO_2_. For the final 48 hours, moDCs were treated with recombinant human IFNα (100 U/ml, Immunotools).

### Generation of Siglec-1-targeting VHHs

VHH’s were generated against human Siglec-1 by immunizing two llamas with cells and recombinant protein in four rounds. The first two rounds consisted of human moDCs, treated with IFNα to upregulate Siglec-1 expression, one round of THP-1 cells expressing Siglec-1 (TSn) [48], and a final round with recombinant human Siglec-1 (Ser20-Gln1641) (R&D Systems, 5197-SL). Peripheral blood was obtained from each llama and PBMCs were isolated after which RNA was isolated and reverse transcribed into cDNA. Genes of the variable domain of heavy chain only antibodies were amplified and ligated into the phagemid vector pPQ81 (QVQ) for generation of phage-display libraries using *E. coli* TG1 cells as previously described [49]. Phage-display selections were performed using 5 µg/mL recombinant human Siglec-1 (Ser20-Gln1641) (R&D Systems, 5197-SL) coated in Maxisorp plates. Blocking was performed with 4% (w/v) skimmed milk in PBS and counter selections were performed using plates coated with human IgG1. Output phages were generated after either chemical or antibody (anti- Siglec-1, clone 7-239, [47]) elution. Afterwards, TG1 cells were infected by output phages and used to generate four 96 well masterplates containing periplasmic extracts which were then screened for binding to recombinant human Siglec-1. A total of 368 clones were screened for binding and subsequently sequenced resulting in 81 family clusters (CDR-HR homology >80%) of which 1-2 clones were selected for VHH production as previously described [49] and further analysis.

### VHH Production

VHH sequences of selected clones were ordered from TWIST Bioscience with an N-terminal PelB sequence and C-terminal Myc and 6xhis tag and subcloned into pET28a+. VHH clones were transformed into LOBSTR-BL21(DE3)-RIL cells (Kerafast, EC1002). Starter cultures were grown in 2 mL Luria’s Broth medium supplemented with 50 μg/mL kanamycin and 2% glucose O/N at 37˚C, shaking at 280 rpm. Next, VHH were expressed by innoculating 500 mL autoinduction medium consisting of terrific broth base including trace elements (Foremedium, AIMTB2010) supplemented with 50 μg/mL kanamycin and grown for 56 hours at 18˚C, shaking at 220 rpm. Cultures were harvested by centrifugation at 4000 rcf for 60 min at 4˚C. Cell pellets were incubated O/N at -20˚C after which they were resuspended in PBS supplemented with 1 mM MgCl_2_ and 5 μg/mL DNase-I and incubated on a rotor O/N at 4˚C. Lysates were clarified by centrifugation at 4000 rcf for 60 min at 4˚C and subsequently filtered through a 0.2 um filter and supplemented with 30 mM imidazole. VHH’s were purified by affinity chromatography using 5 mL His-Trap HP columns (Cytiva) and afterwards imidazole was removed by gel filtration with a HiPrep™ 26/10 column (Cytiva).

### Human Siglec-1 ELISA

His-tagged recombinant human Siglec-1 (Ser20-Gln1641) (R&D Systems, 5197-SL) or Siglec-1 domain 1 (Ser20-Thr136) or D1-2 (Ser20-Glu233) were coated with a concentration of 5 µg/mL in coating buffer (0,2 M NaHCO_3_, pH=9,7) in Nunc MaxiSorp ELISA plates (Thermo Fisher Scientific) O/N at 4°C. Plates were subsequently blocked with carbo free blocking buffer (Vector labs, SP-5040-125) for 30 minutes at 37°C. Next, VHH and antibody incubations were performed in carbo free blocking buffer 1 hour at RT. VHH were detected with anti-his horseradish peroxide (HRP) conjugated antibodies (Biolegend). Anti-Siglec-1 antibodies with anti-mouse-HRP (Agilent, P0260). After each step wells were washed with PBS supplemented with Tween-20 (0.05% w/v). ELISA was developed by the addition of TMB substrate (ThermoFisher, N301), the reaction was stopped by the addition of 0.3 M H_2_SO_4_ and absorbance was measured at 450 nm using a microplate spectrophotometer.

### VHH conjugation with Alexa Fluor 647

VHH’s were expressed and purified with a C-terminal 6xhis-tag separated by a rigid-linker (SPSTPPTPSPSTPP) from a cysteine (AAK**<u>C</u>**KAA) used for maleimide conjugation (Massa et a., 2014). The VHH were subsequently reduced by adding 50 mM TCEP (Sigma Aldrich) at a molar ratio of TCEP:VHH of 3:1 for 2 hours at room temperature. Next, TCEP was removed by buffer exchange using a PD10 desalting column and Alexa Fluor™ 647 C_2_ Maleimide dissolved in DMSO was added to the VHH and incubated O/N at 4°C on a roller in the dark. Next, free dye was removed with a PD10 desalting column. VHH were then concentrated using Vivaspin tubes (Sigma Aaldrich, Z629464) with a molecular weight cut-off of 3 kDa.

### VHH and antibody binding to cells

Cells were first pre-incubated with human Fc Block (BD Biosciences), True-Stain Monocyte Blocker (Biolegend) and viability dyes (Fixable viability dye eFluor 780, eBioscience, or Live Dead Blue, Life Technologies) in PBS at 4°C for 10 min. For cells expressing Siglec-1-halotag, cells were stained with cell permeable halotag ligand (CA-MaP555, Tenova Pharmaceuticals) on ice for 30 minutes followed by five washing steps with PBS supplemented with BSA (0.5% w/v) (PBA). Next, cells were incubated at 4°C with either VHH or antibodies for 30 minutes at 4°C followed by washing with PBA after each antibody incubation step. VHH were detected with anti-Myc-tag antibody (Cell Signaling) or anti-his tag-AF647 antibody (Biolegend). In the epitope binning experiments VHH were labelled with AF647 dyes. Cells were fixed with 2% paraformaldehyde (PFA) for 10 min at 4°C. Cells were stored at 4°C up to 1 week until acquisition on Attune (Life Technologies), BD LSR Fortessa (BD Bioscience), or Aurora spectral flow cytometer (Cytek) and analyzed on FlowJo software (Tree Star) or OMIQ (Dotmatics). In the case of pLent6.3-Siglec-1-halotag transfected HEK293T cells, successfully transfected cells were identified by halotag ligand Ca-MaP555 positive signal. VHH binding curves were analyzed with GraphPad Prism 10.6 and fitted with four parameter dose-response curve.

### Production of recombinant human Siglec-1 domain 1 and 2

Siglec-1 gene fragments encoding Siclec-1-D1 (Ser20-Glu138) and Siglec-1 D1D2 1 (Ser20-233Gln) were cloned into pFastBacNKI-GP67-LIC-3C-2xStrepII-10xhis, using ligation independent cloning[50]. Proteins are expressed with an N-terminal GP67 signal peptide and a C-terminal TwinstrepII-10xhis tag. Plasmids were transformed into EMBACY cells and bacmid DNA was isolated from three individual positive (white) colonies, according to the Bac-to-Bac manual (Thermofisher Scientific). Sf9 insect cells were plated in 6-well plates and the cells were transfected with bacmid DNA. After three days at 28 °C, the medium containing secreted virus particles was harvested (P0 virus) and added to 25 ml of sf9 suspension cultures. After three days, cells were monitored for growth, cell diameter and YFP expression to confirm virus production. Cells were removed by centrifugation (4 minutes at 900 x g) and medium was collected (P1 virus stock).

For protein expression, sf9 cells were infected with P1 virus (1 ml P1 virus added to 500 ml 1 x 10^6^ cells/ml sf9 cells). After three days, cells were removed by centrifugation (15 minutes at 900 x g) and medium was collected. The pH of the medium was adjusted to pH 8.0 by addition of 50 ml Tris-HCl pH 8.0 to 500 ml solution and precipitates were removed by centrifugation (10 minutes at 8,000 x g). The solution was passed over 1.5 ml of Nickel Indigo beads (Cube Biotech). Beads were washed with 15 ml 25 mM HEPES pH 7.5, 200 mM NacL, 20 mM imidazole and protein was eluted using the same buffer including 300 mM imidazole. Elution fractions were pooled and passed over 1 ml of Streptactin XT beads (IBA Lifesciences). Beads were washed with 25 mM HEPES pH 7.5, 200 mM NacL before Siglec-1 proteins were eluted in the same buffer, supplemented with 50 mM biotin. Purified proteins were concentrated in Vivaspin concentrators (VIVAproducts) and dialyzed against 1 liter of 25 mM Hepes pH 7.5, 200 mM Nacl. Protein aliquots were snap frozen in liquid nitrogen and stored at -80 °C.

### Construction of Siglec-1-halotag expression vector

Siglec-1 WT and R97A (R116A when signal peptide is included) receptor mutant were designed with a c-terminal halotag separated by a small flexible linker (GGGGSGGGGS). Siglec-1 DNA inserts were ordered from Twist Bioscience in two parts: part-1 (Siglec-1: 1-3371 bp) had a 5’ BamHI and 3’ XbaI restriction site, part-2 (Siglec-1: 3372-5127 bp + halotag) had a 5’ XbaI and 3’ BstBI restriction site. A pLenti-6.3 expression vector with a CMV promoter transcribing a 5’ BamHI and 3’ XbaI flanked irrelevant gene with downstream 3’ BstBI restriction site. The irrelevant gene was digested with BamHI and XbaI (FastDigest, Thermo Fisher) before digestion and ligation part-1 into the vector. Next, the pLenti-6.3-Siglec-1-Part-1 vector was digested with XbaI and BstBI and Siglec-1-halotag was digested and ligated into the vector to generate a pLenti-6.3-Siglec-1-halotag vector containing WT or R97A mutant Siglec-1. Vector construction was verified by wholeplasmid sequencing (Eurofins).

### Siglec-1-halotag expression

HEK293T cells were seeded in T75 plates at 3 million cells per flask in DMEM (10% FCS, 1% penicillin and streptomycin) and incubated O/N at 37°C and 5% CO_2_. The next day medium was refreshed and cells were transfected with 10 µg receptor DNA using the PEI method in a 1:6 ratio of DNA to PEI (Polysciences Inc) and incubated O/N at 37°C and 5% CO_2_. The following day medium was refreshed and cells were incubated another 48 hours at 37°C and 5% CO_2_. Next, cells were collected by scraping and used for flow cytometry analysis as described above.

### Surface plasmon resonance binding experiments

The binding affinities of VHHs to Siglec-1 were determined using surface plasmon resonance (SPR) on a Biacore 1S+ instrument (Cytiva Life Sciences). Siglec-1-FL was immobilized on a CM5 sensor chip (Cytiva Life Sciences) via amine coupling in 10 mM glycine buffer, pH 5.0. Serial twofold dilutions of VHHs were injected over the chip surface in HBS-P+ buffer (10 mM HEPES, pH 7.4; 150 mM NaCl; 0.05% v/v Surfactant P20; Cytiva Life Sciences) using a single-cycle injection procedure. Binding responses were processed by double referencing, involving subtraction of both the reference surface and buffer blank responses. Data were analyzed using Biacore Insight Software (Cytiva Life Sciences). Binding data for 1B5 and 2C2 were analyzed using a 1:1 kinetic binding model fit. Due to its very fast dissociation rate, which approached the experimental detection limit, the binding data for 1C1 were analyzed using an equilibrium binding model. Figures were prepared using GraphPad Prism (GraphPad Software, LLC).

### Alpha Fold VHH binding prediction

Alpha Fold3 was used to predict VHH binding to human Siglec-1. The input sequence of human Siglec-1 (Uniprot Q9BZZ2) consisted of the first domain (Ser20-Glu138) for modeling interactions with VHH clone 2C2 and the first two domains (Ser20-233Gln) for modeling interactions with VHH clones 1B5 and 1C1. Models were analyzed using UCSF Chimera-X (Meng EC, Goddard TD, Pettersen EF, Couch GS, Pearson ZJ, Morris JH, Ferrin TE. Protein Sci. 2023 Nov;32(11):e4792.). Coloring of the structures according to the pLDDT scores was done according to the color key shown in the figures (<50, 50-70, >90).

### Liposome binding

Siglec-1-targeting liposomes were generated as previously described[51]. Liposome binding experiments were performed with BwSn cells. Cells were pre-incubated with VHH or antibodies as indicated before being incubated with ganglioside-liposomes (100 µM) for 30 min at 4°C. In the case of neuraminidase treatment, cells were washed three times with PBS prior to incubation with 0.4 U/mL of Neuraminidase from Clostridium perfringens (Roche, Cat. No.: 11585886001) or as control PBS, for 1 hour at 37°C, shaking at 700 rpm. Cells were subsequently washed three times with carbo free blocking buffer (Vector Labs, Cat. No.: SP-5040-125) before VHH/liposome binding. De-sialylation was evaluated by loss of biotinylated Maackia Amurensis Lectin II (MAL II) (Vector Labs, Cat. No.: B-1265-1) binding. Cells were then fixed and acquired on flow cytometry. Liposome binding curves were fitted with log(inhibitor) vs. response -Variable slope in GraphPad Prism 10.6 to determine VHH IC_50_ values.

### Campylobacter jejuni FITC labelling

In this study GBS associated *C. jejuni* strains GB14 (ganglioside mimic GM1a, LOS class C), GB23 (ganglioside mimic GM2, LOS class A) and GB31 (ganglioside mimic GD1a, GM1a, LOS class A) were used (previously described by Heikema et al., 2010). The *C. jejuni* strains originate from a GBS–related strain collection of the Erasmus MC, Rotterdam, the Netherlands. Freshly cultured *C. jejuni* were harvested in PBS and incubated with fluorescein isothiocynanate (FITC) with shaking for 60 min at RT in the dark. Bacteria were then washed three times in PBS before being resuspended in PBS supplemented with 2 mM MgCl_2_. Afterwards, *C. jejuni* was heat inactivated by 45 minute incubation at 56°C in the dark and subsequently washed three times with PBS after which bacterial solutions were transferred to storage medium (PBS + glycerol) and stored at -80°C. Thawed bacteria were washed two times in PBS before being used in binding experiments.

### Campylobacter jejuni binding

A cell to bacteria ratio of 1:10 or 1:100 was used in binding experiments. Cells were pre-incubated with VHH or antibodies where indicated for 30 minutes at 4°C before being incubated with FITC labelled *C. jejuni* for 2 hours at 37°C for binding. After which cells were washed once with RPMI supplemented with 1% FCS and then stained for 10 min with Fixable viability dye eFluor 780 (eBioscience) for 10 minutes at 4°C. Cells were then washed with PBA before being fixed with 2% paraformaldehyde for 10 minutes in the dark at 4°C. Binding was then measured by flow cytometry. In experiments where cytokine secretion was measured, moDCs were pre-incubated with VHH and antibodies before a six hour incubation with FITC labelled *C. jejuni*. Next, cells were pelleted by centrifugation and the supernatant was subjected to an IL-6 ELISA (Biolegend, Cat. No.: 430501). In the experiments were peripheral blood mononuclear cells (PBMCs) were required they were isolated from heparinized blood by density gradient centrifugation (Lymphoprep; Axis-Shield PoC AS). Following PBS washes, cells were further processed for *C. jejuni* binding and flow cytometry, as described above. *C. jejuni* binding curves were fitted with log(inhibitor) vs. response -Variable slope in GraphPad Prism 10.6 to determine VHH IC_50_ values.

### SARS-CoV-2 recombinant protein binding

BWSn cells were stained for 10 minutes with fixable viability dye eFluor 780 (eBioscience) at 4°C. Cells were then incubated with VHH or 7-239 antibody for 30 min at 4°C. Next, recombinant spike protein S1+S2 trimer (Sino Biological, 40589-V49H-B) was added and incubated with cells for 30 min at 4°C. Afterwards, cells were washed two times with PBA before spike protein was detected with streptavidin-PE (Agilent, PJRS25-1). Cells were then washed two times with PBA followed by fixation with 2% PFA for 20 minutes. Cells were then analyzed by flow cytometry.

### SARS-CoV-2 virus production

The following reagent was obtained from Dr. Maria R. Capobianchi through BEI Resources, NIAID, NIH: SARS-related coronavirus 2, Isolate Italy-INMI1, NR-52284, originally isolated in January 2020 in Rome, Italy. VeroE6 cells (ATCC CRL-1586) were inoculated with the SARS-CoV-2 isolate and used for reproduction of virus stocks, as described previously [52].

### SARS-CoV-2 virus binding

THP-1 and TSn cell or moDCs were incubated with the SARS-CoV-2 isolate (hCoV-19/Italy; TCID50 10^4^ for 4 h at 37°C. After 4 h cells were washed extensively to remove unbound virus. Cells were lysed and cellular and viral RNA was isolated using the QIAamp Viral RNA Mini Kit (Qiagen, Germany) according to the manufacturer’s protocol. cDNA was synthesized with the M-MLV reverse-transcriptase kit (Promega, USA) and diluted 1:5 before further application. PCR amplification was performed in the presence of SYBR green (Thermo Fisher Scientific, USA) in a 7500 Fast Realtime PCR System (Applied Biosystems, USA). The normalized amount of target mRNA was calculated from the Ct values obtained for both target and household mRNA with the equation Nt = 2Ct ^(GAPDH)-Ct(target)^. The following primers were used: GAPDH, forward primer (CCATGTTCGTCATGGGTGTG), reverse primer (GGTGCTAAGCAGTTGGTGGTG). ORF1b, forward primer (TGGGGTTTTACAGGTAACCT), reverse primer (AACACGCTTAACAAAGCACTC)[53].

### Statistics

Statistical analyses were performed using one-way ANOVA with Dunnett’s multiple correction or uncorrected Fishers LSD t-test, Welch’s unpaired t-test were performed with Graphpad Prism 10.6 (Graphpad software).

## Results

### Immunization strategy and selection of Siglec-1 binding VHH

To generate human Siglec-1-specific VHH two llamas were immunized twice with human moDC treated with type I interferon (IFN-I) to upregulate Siglec-1, once with THP-1 cells overexpressing Siglec-1 (TSn) and finally a single round with recombinant Siglec-1 consisting of all extracellular domains (D1-17, Ser20-Gln1641) (Fig. S1A). Phage-display libraries were generated for panning against recombinant human Siglec-1 using a phage ELISA. Next, clones were selected for VHH production and screened as periplasmic extracts for binding to cellular and recombinant Siglec-1 by flow cytometry and ELISA, respectively (Fig. S1B). VHH clones were subdivided into 86 VHH families based on their CDR3 regions (similarity ≥ 80%). A selection of 22 clones were produced as recombinant protein and tested for binding to Siglec-1 expressing BW517 cells (BWSn) by flow cytometry, of which only 18 clones showed binding to Siglec-1 (Fig. 1B). Next, VHH clones 1B5, 1C1 and 2C2 were selected for further characterization. Half-maximal binding concentrations were determined for binding to BWSn cells. 1B5 and 2C2 displayed half-maximal binding concentrations in the low nanomolar range of 10 ± 3 nM and 15 ± 9 nM, similar to the Siglec-1-specific monoclonal antibody (mAb) 7-239 3 ± 1 nM, while that of 1C1 was in the higher nanomolar range 170 ± 95 nM (Fig. 1C). In conclusion, 18 VHH clones were identified that bind to human Siglec-1 and three clones 1B5, 1C1 and 2C2 were selected for further characterization.

**Fig. 1.**
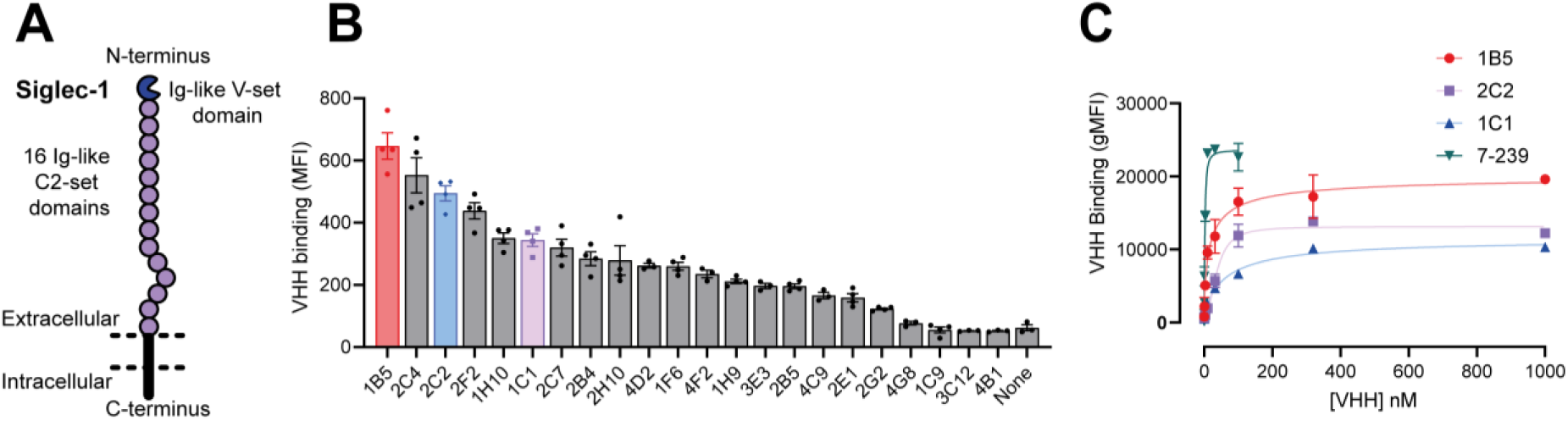
Selection of Siglec-1-binding VHH. **A)** Schematic overview of Siglec-1. **B)** VHH binding (500 nM) to Siglec-1 expressing BW5147 cells (BWSn) detected by flow cytometry. Data is shown as mean fluorescent intensity. **C)** Concentration binding curves of selected VHH clones 1B5, 2C2 and 1C1 and antibody 7-239 binding to BWSn cells. B) Data is shown as mean ± SEM of n=3 experiments. C) A representative figure is shown of n=3 experiments. Data is shown as mean ± SD.

### Epitope mapping of Siglec-1 binding VHH

Siglec-1 is a large type 1 membrane receptor with 17 extracellular domains of which the terminal V-set domain 1 is responsible for binding to sialic acids. Ligand binding can be blocked by mAb 7D2 and 7-239 that are known to recognize different epitopes (Fig. 1A)[27, 47, 54]. First, we performed blocking assays with VHH and mAb 7D2/7-239. Pre-incubation of cells with 7D2 blocked binding of all three VHH and in the reverse set up all three VHH blocked 7D2, which suggests that all three VHH bind to or near the terminal domain (Fig. 2A, S2A). In contrast, pre-incubating cells with 7-239 significantly increased the binding of VHH clones 1B5 and 1C1, whereas 2C2 binding was efficiently blocked (Fig. 2B). In the reverse set up, 7-239 binding was significantly increased by pre-incubation with VHH clone 1B5 but not affected by 1C1 or 2C2 pre-incubation (Fig. 2B, S2A). Next, displacement experiments were performed in which BWSn cells were first pre-incubated with AF647-labelled VHH and then unlabeled VHH to determine their capacity to displace Siglec-1 bound labelled VHH. 1B5 and 1C1 could partially displace each other while 2C2 binding was unaffected (Fig. 2C, S2B). Conversely, 2C2 was not capable of displacing 1B5 or 1C1 (Fig. 2C, S2B). Taken together, these results imply that 1B5 and 1C1 have a similar epitope that does not overlap with 2C2, while the epitope of 2C2 overlaps with 7-239. In addition, the binding of 1B5 or 1C1 together with 7-239 appeared to be mutually enhancing.

**Fig. 2.**
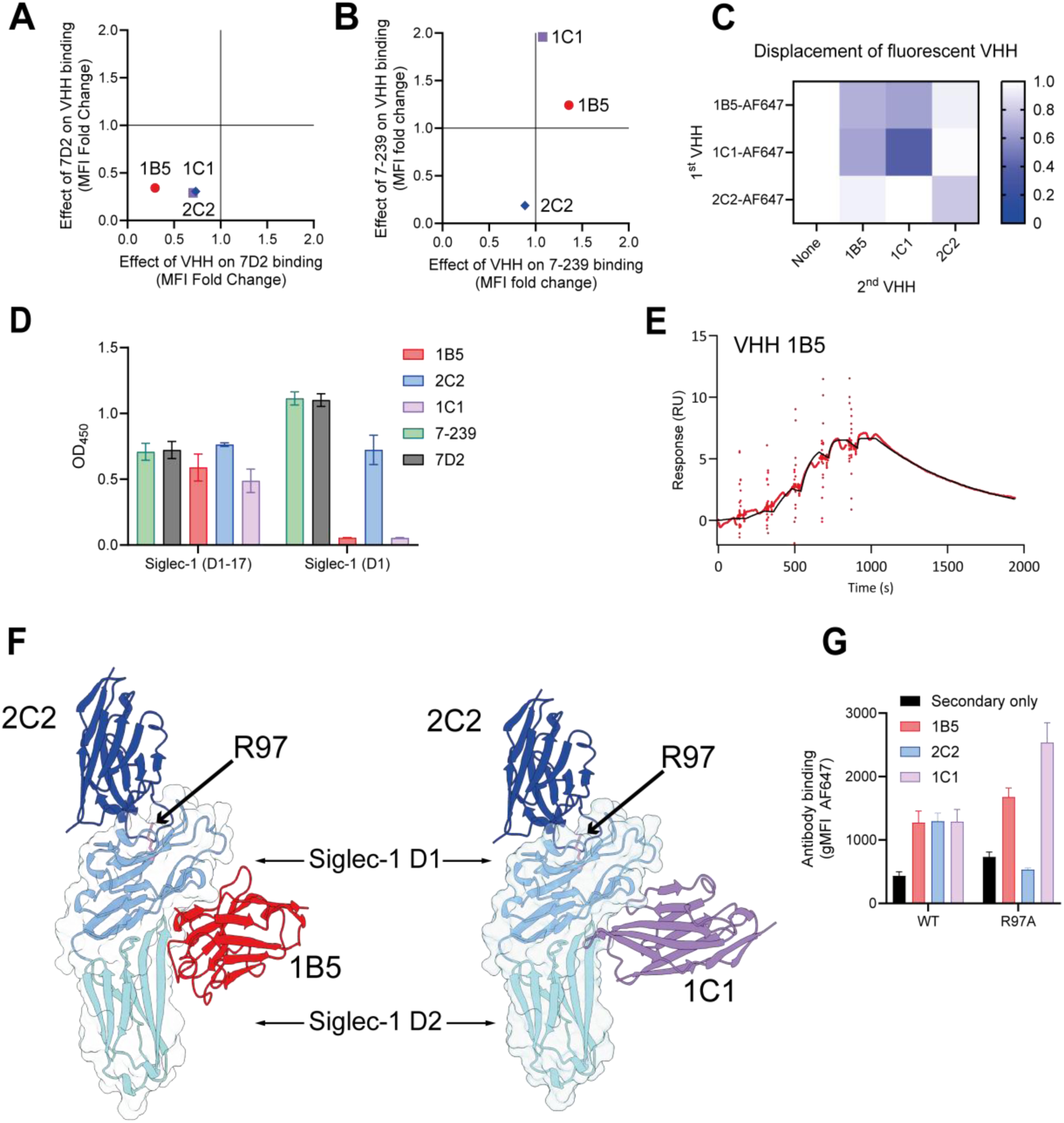
Epitope mapping of Siglec-1 binding VHH. A-B) Effect of pre-incubation with antibody 7D2 or 7-239 on VHH binding to BWSn (y-axis) or the reverse, in which cells were pre-incubated with VHH followed by 7D2 or 7-239 binding (x-axis). Binding was detected by flow cytometry and shown as fold change of mean fluorescent intensity data after normalization to a no antibody/VHH control for n=4 experiments. **C)** Binding data of AF647 labelled VHH after displacement with non-labelled VHH to BWSn cells. Data is normalized to the no competing VHH condition of n=3 experiments. **D)** ELISA binding data of VHH and antibodies to recombinant Siglec-1 (Ser20-Gln-1641) and Siglec-1 domain 1 (Ser20-Thr136). Data is shown as mean ± SEM of n=3 experiments. **E)** Binding of VHH 1B5 to immobilized recombinant Siglec-1 (Ser20-Gln-1641) as measured by surface plasmon resonance. Data was fitted using a 1:1 kinetic binding model fit. **F)** Alphafold3 model of VHH clone 2C2 (blue) combined with either 1B5 (red) or 1C1 (purple) binding to Siglec-1 domain 1-2. Residue R97 of Siglec-1 is indicated by an arrow. **G)** VHH and antibody binding to HEK293T cells expressing Siglec-1 WT or R97A mutant receptor. Data is shown as mean ± SD of a representative figure of n=3 experiments.

Considering that mAb 7D2 blocks ligand binding and the binding of all three VHH was inhibited by 7D2, we hypothesized that the epitopes of all VHH are located on or near the terminal Ig-like V-set domain 1 of Siglec-1 (Siglec-1-D1)[51]. We therefore generated recombinant Siglec-1-D1 and tested binding of our VHH in an ELISA using recombinant Siglec-1 D1 or full length Siglec-1-D1-17. VHH 2C2 and mAb 7-239 and 7D2 bound to both Siglec-1-D1 and Siglec-1-D1-17, indicating that they have epitopes in D1 (Fig. 2D). Interestingly, 1B5 and 1C1 did not bind to Siglec-1-D1, but only to Siglec-1-D1-17. Since 1B5 and 1C1 were also displaced by 7D2 that binds to D1, this suggested that they might have an epitope on the interface between Siglec-1-D1 and D2. We next determined the affinities of our VHH using Siglec-1-D1-17 in surface plasmon resonance (SPR) measurements (Fig. 2E, S2C,D). VHH clones 1B5 and 2C2 bound with a high affinity (0.6 and 7.4 nM), while 1C1 displayed a lower affinity (21 nM).

Finally, we modeled the binding epitopes of our VHH using Alphafold3 with either Siglec-1-D1 or Siglec-1-D1-2 as input. No crystal structure currently exists for human Siglec-1, but the Ig-like V-set domain (D1) of mouse Siglec-1 has been crystallized [55]. The Alphafold3 fold of human Siglec-1-D1 was structurally very similar to mouse Siglec-1 with a high predicted template modeling score (pTM; 0.9) (Fig. S2E,F). VHH clone 2C2 was preferentially modeled on the ligand binding site of Siglec-1-D1 in a head-on orientation [56]. The Siglec-1-D1/2C2 fold had a high pTM score of 0.86 and high interface predicted template modeling (ipTM=0.8) score and overall confident predicted local distance difference test score (pLDDT>0.7) with the exception of CDR3 (Low (70 > plDDT > 50)) (Fig. 2F, S2G, H). VHH clones 1B5 and 1C1 were modeled onto the interface between Siglec-1-D1 and 2, opposite the ligand binding domain, with a high modelling confidence for both 1B5 (pTM=0.86, ipTM=0.88) and 1C1 (pTM=0.80, ipTM=0.85) (Fig. 2F, S2G, H). 1B5 was predicted to have a side-on interaction whereas 1C1 assumed a head-on binding pose similar to 2C2. Interestingly, 2C2 was modeled onto R97 of Siglec-1-D1, which is known to be essential for ligand binding as mutation of this residue (R97A) abolishes Siglec-1 ligand interactions [57]. In the Alphafold3 model, the mutation R97A decreased output confidence significantly (pTM=0.27, ipTM=0.6). We therefore tested the binding of our VHH to HEK293T cells expressing either WT or R97A Siglec-1 by flow cytometry. 1B5 and 1C1 could bind to both Siglec-1 forms, whereas 2C2 could only bind to WT Siglec-1 but not the R97A mutant, supporting the Alphafold3 prediction (Fig. 2G). In conclusion, VHH 2C2 binds specifically to the ligand binding site of Siglec-1, whereas VHH 1B5 and 1C1 have predicted epitopes on the interface between D1 and D2.

### Siglec-1 binding VHH ortho- and allosterically modulate ligand binding

Having established that our VHH bind to (2C2) or away from (1B5/1C1) the ligand binding domain, we subsequently investigated the effect of our VHH on ligand binding. Siglec-1 is a receptor for gangliosides present on the membrane of viruses or ganglioside-like LOS structures on bacteria [13, 31]. Ganglioside-containing liposomes bind to Siglec-1 and mimic these interactions (Fig. 3A). To address whether our selected VHHs can interfere with Siglec-1/ganglioside interactions, we pre-incubated Siglec-1 expressing cells with various concentrations of VHH or blocking mAb 7-239 [58], washed and incubated with DiD-labeled GM3 liposomes (Fig. 3A, B, Fig. S3A). Clone 2C2 completely blocked liposome binding (IC_50_: 26.3 ± 2.5 nM), similar to mAb 7-239, and since it binds to the ligand binding site, it appears to be an orthosteric modulator (Fig. 3B, S3B). Unexpectedly, clones 1B5 and 1C1 displayed a concentration-dependent enhancement of GM3 liposome binding (Fig. 3B). To determine whether the blocking or enhancing effect of the VHHs on Siglec-1 ligand interactions was specific to GM3 or a more general effect for Siglec-1-binding gangliosides, GM1, GM3, GD1a and GD3 liposomes were tested. Pre-incubation with 500 nM 2C2 blocked liposome binding to Siglec-1 regardless of the ganglioside ligand used to a similar degree as 100 nM of 7-239 (Fig. 3C). 1B5 significantly enhanced ligand binding independent of the ganglioside tested, while 1C1 showed a similar, although non-significant, trend. Since both 1B5 and 1C1 do not bind at the ligand binding site, these two VHH appear to affect ligand binding allosterically (Fig. 3C). In conclusion, VHH 1B5, 1C1 and 2C2 modulate ligand binding by either enhancing or blocking interactions between ganglioside-containing liposomes and Siglec-1.

**Fig. 3.**
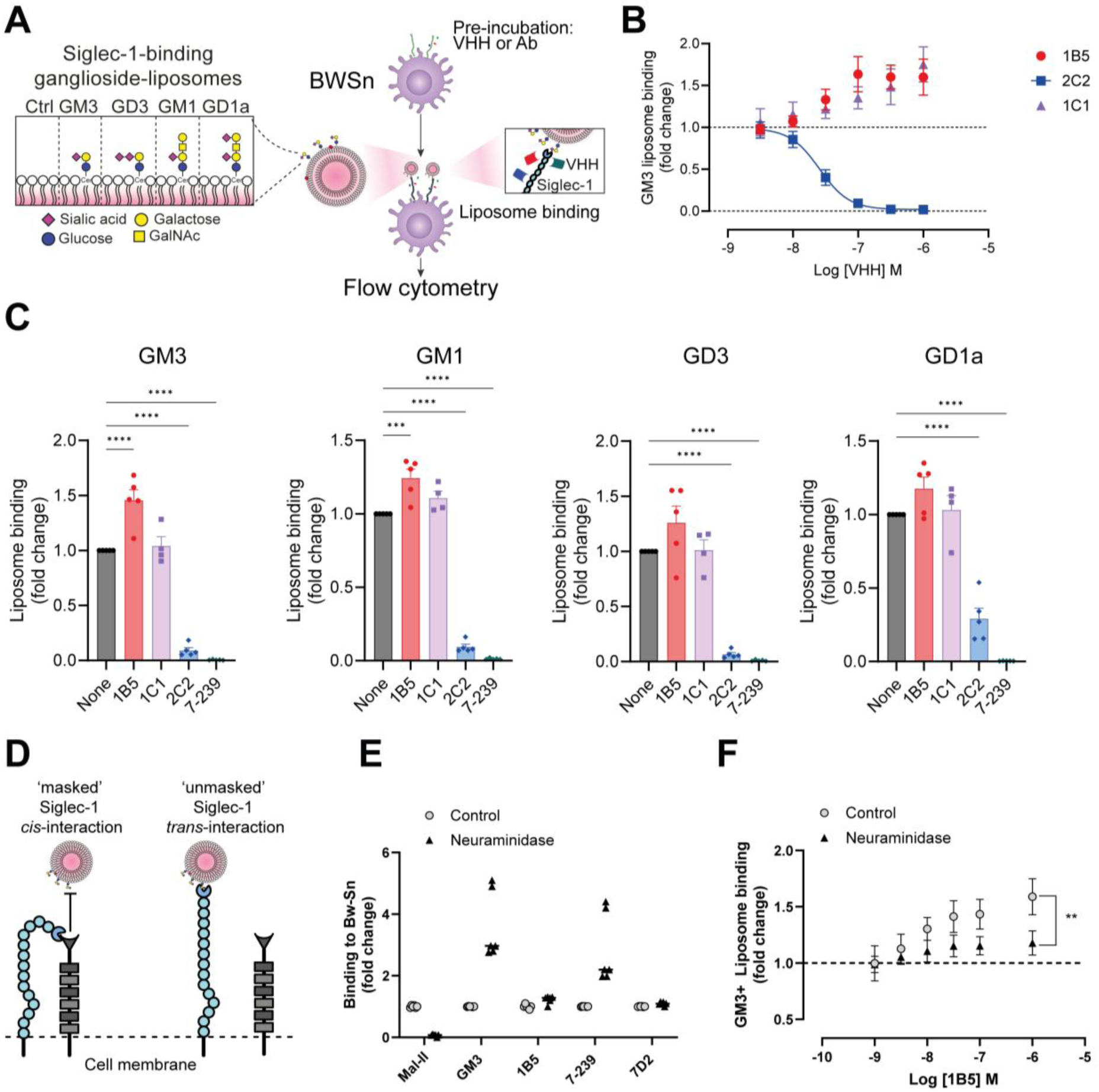
VHH mediated modulation of ganglioside liposome binding to Siglec-1. A) Schematic overview of experiments to determine effect of VHH on ganglioside liposome binding. **B)** GM3 liposome binding after pre-incubation of BWSn cells with various concentrations of VHH. **C)** Ganglioside liposomes binding to BWSn cells after pre-incubation with VHH or antibody 7-239. **D)** Cartoon of Siglec-1 *cis* and *trans* interactions. **E)** Binding of lectin MALl-II, GM3 liposomes, VHH 1B5, antibodies 7-239 and 7D2 to BWSn cells that were either treated with neuraminidase or a buffer control. **F)** GM3 liposome binding after pre-incubation of BWSn cells with various concentrations of VHH 1B5. Cells were treated prior to binding with neuraminidase or a buffer control. Liposome binding detected by flow cytometry. Fold change data is shown of liposome DiD signal normalized to a no VHH control. Data is shown as mean ± SEM of B, E, F) n=3 or C) n=5 experiments (t-test:\*\**p<0.01*, one-way ANOVA: \*\*\*\**p<0.0001*).

### VHH 1B5 potentially modulates Siglec-1 *cis-*interactions

Siglec-1 is known to engage ligands in both *trans* and *cis-*interactions whereby *cis*-interactions ‘mask’ the receptor for binding to soluble ligands and ligands on other cells (Fig. 3D). Therefore, we speculated that the enhancement in liposome binding achieved by pre-incubating with VHH 1B5 and 1C1 could partially be the result of the VHH modulating *cis*-interactions. To test this hypothesis, we pre-treated BWSn cells with neuraminidase to remove sialic acid from the surface of the cells resulting in unmasking of Siglec-1 from *cis*-interactions. Successful desialylation of surface exposed sialic acid on BWSn cell was confirmed by a significant reduction in binding of MAL-II, a plant lectin that specifically engages sialic acid in an (α-2,3) linkage on O-glycans exposed on the cell surface (Fig. 3E). Binding of GM3 liposomes and mAb 7-239 increased significantly when cells were treated with neuraminidase, but VHH 1B5 and mAb 7D2 binding was only enhanced to a small extent (Fig. 3E). Next, cells treated with neuraminidase were pre-incubated with VHH 1B5 and subsequently incubated with DiD-labeled GM3 liposomes. Overall, GM3 liposome binding was significantly enhanced by neuraminidase, but the VHH 1B5 mediated effect was significantly reduced for the neuraminidase-treated cells when compared to the buffer only control (Fig. 3F). Taken together, these results suggest that the ligand binding enhancing effect of VHH 1B5 could be mediated through modulation of Siglec-1 *cis-* interactions.

### VHH 2C2 blocks Siglec-1-mediated *C. jejuni* binding

Siglec-1 binds pathogens such as the gram-negative bacterium *C. jejuni* that expresses LOS structures with sialylated moieties mimicking Siglec-1-binding gangliosides on its outer membrane. To investigate the modulating effect of our VHHs on the binding of *C. jejuni* to Siglec-1 expressing cells, we incubated TSn with heat-inactivated FITC-labeled *C. jejuni* strains expressing various ganglioside mimics (GB14 = GM1a, GB23 = GM2, GB31 = GM1a/GD1a) [39]. All three *C. jejuni* strains bound to TSn cells but not to the parental cell line THP-1 (Fig. S4B). Dose titration curves demonstrated a strong blocking effect of 2C2 and an enhancing effect of 1B5 on *C. jejuni* binding for all three strains tested (Fig. 4A). The binding of the *C. jejuni* strains to TSn cells was also visualized by fluorescent microscopy (Fig. 4B). A clear association with the cell membrane is observed in the untreated condition or in the presence of 1B5, whereas 2C2 and 7-239 blocked all three *C. jejuni* strains from binding TSn cells (Fig. 4B, S4C).

**Fig. 4.**
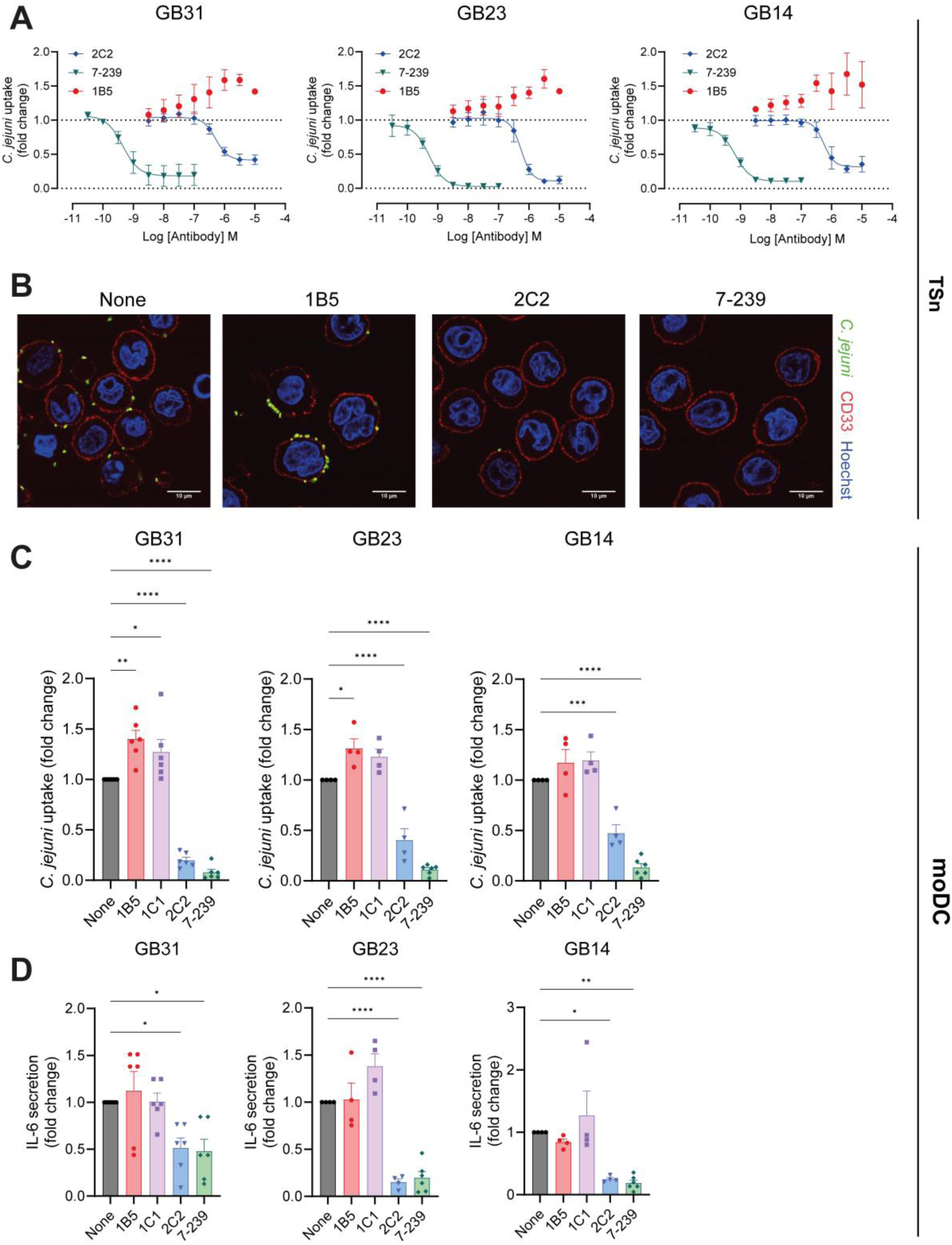
Siglec-1-binding VHH modulate *C. jejuni* binding and uptake. **A)** Binding data of FITC-labeled *C. jejuni* strains GB31, GB23 and GB14 to Siglec-1 expressing THP1 cells (TSn) as determined by flow cytometry. TSn cells were pre-incubated with various concentrations of VHH or antibody 7-239 at 4°C before incubation with *C. jejuni* Mean fluorescent intensity binding data is normalized to a no VHH/antibody control. Data is shown as mean ± SEM of n=3 experiments. **B)** Representative fluorescent microscopy images of FITC-labelled GB31 *C. jejuni* binding to TSn cells pre-incubated with 7-239 or VHH. TSn membranes detected with anti-CD33-AF647 antibody. Nucleus is stained with Hoechst. **C)** Uptake of *C. jejuni* strains GB31, GB23 and GB14 by moDCs pre-incubated with VHH or antibody as determined by flow cytometry. Normalized data is shown as fold change over no VHH control of 4 to 6 donors. (one way ANOVA: \*\**p<0.01, ***p<0.001, ****p<0.0001*) **D)** IL-6 secretion by moDCs, pre-incubated with VHH or antibody, followed by a 6 hour incubation with *C. jejuni* strains GB31, GB23 and GB14 at 37°C. IL-6 amounts were determined by ELISA. Normalized data is shown as fold change over no VHH control of 4 to 6 donors (one way ANOVA: \**p<0.05, **p<0.01*, *\*\*\*\*p<0.0001*).

Next we assessed the effects of the three VHH on binding of *C. jejuni* strains GB14, 23, and 31 by Siglec-1^+^ moDCs. Although moDCs express multiple Siglec receptors, binding of all strains by moDCs was Siglec-1 dependent as it could be blocked by pre-incubation with 7-239 similar to what was observed for macrophages [39] (Fig. 4C). Pre-incubation with 500 nM of 2C2 significantly blocked *C. jejuni* binding regardless of the strain, while 1B5 and 1C1 demonstrated an enhanced binding of all strains (Fig. 4C, S4D). All three *C. jejuni* strains tested induced production of IL-6, which could, to a large extent be abrogated by pre-incubation with 7-239 and 2C2, whereas VHH alone could not activate moDCs (Fig. 4D, S4E). In conclusion, these data reveal that our VHH 1B5 and 1C1 enhance the binding function of Siglec-1, while 2C2 is capable of blocking *C. jejuni* binding and subsequent activation of moDCs.

### Siglec-1-binding VHH bind to primary cells

Previously, we have shown that circulating immune cells in the blood, Axl^+^ Siglec-6^+^ DC cells (ASDCs) (also known as Axl^+^ DC, pre-DC or transitional DC) express Siglec-1 constitutively and can bind ganglioside liposomes [58]. To determine VHH binding to human Siglec-1^+^ primary cells, PBMCs were isolated from healthy donors, and VHH binding was assessed *ex vivo*. Uniform Manifold Approximation and Projection (UMAP) dimensionality reduction analysis was performed on HLA-DR^+^ Lin^-^ DCs using markers for DC lineages: CD14, CD163, CD123, CD11c, CD1c, CD141, Axl, Siglec-6, and Siglec-1. Various DC subsets, including DC1, DC2, DC3, pDC, and Axl^+^ DC cells, were identified with conventional sequential gating strategy and overlaid on the UMAP (Fig. 5A, S5A). Siglec-1 was shown to be predominantly expressed by Axl^+^ DCs and 1B5, 1C1 and 2C2 were shown to bind to this cluster specifically (Fig. 5A-B).

**Fig. 5.**
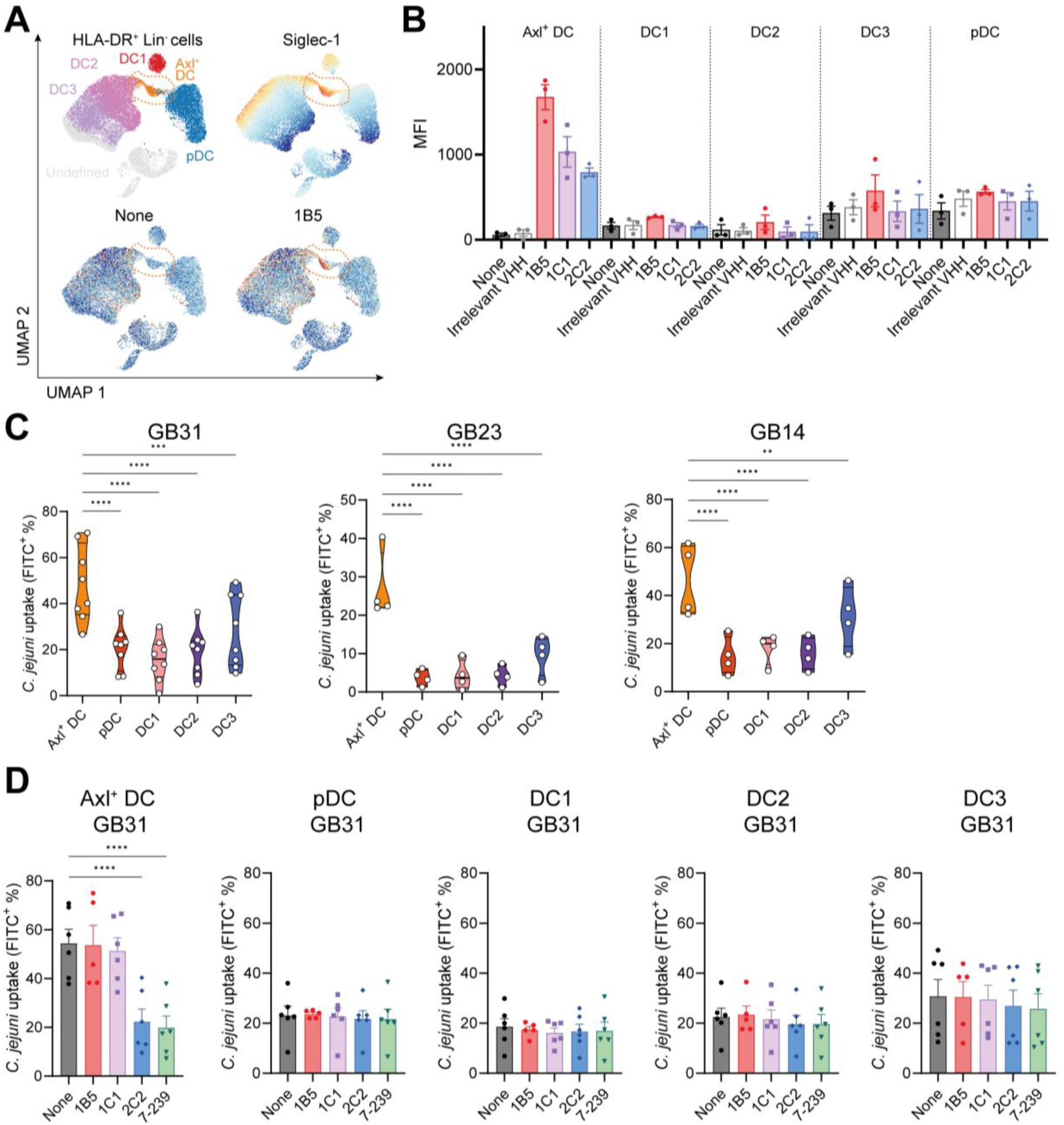
VHH bind to Siglec-1-expressing DCs and VHH 2C2 blocks *C. jejuni* uptake. **A)** High dimensionality reduction analysis by Uniform Manifold Approximation and Projection (UMAP) of HLA-DR^+^ Lin(CD3/CD19/CD56/CD88)^-^ dendritic cells (DCs) using markers: CD123, CD11c, CD1c, CD141, Axl, Siglec-6, and Siglec-1. Identification of DC subsets DC1, DC2, DC3, pDC, and Axl^+^ DC cells was done by conventional sequential gating. Binding of Siglec-1 specific VHH is restricted to cells expressing Siglec-1 on UMAP. **B)** Binding of VHH to DCs as determined by flow cytometry. Data is shown as mean fluorescent intensity (MFI) of at 3 donors. **C)** Percentages of FITC+ cells after incubation with FITC-labelled *C. jejuni* strains GB31, 23 and 14 as determined by flow cytometry. Data is shown for 4 to 8 donors. (one way ANOVA: \*\**p<0.01, ***p<0.001, ****p<0.0001*). **D)** Percentages of FITC+ cells after pre-incubation with VHH or antibody 7-239 followed by incubation with FITC labelled *C. jejuni* strains GB31, 23 & 14 as determined by flow cytometry. Data is shown for 6 donors. (one way ANOVA: *\*\*\*\*p<0.0001*).

### Axl^+^ Siglec-6^+^ DC cells take up *C. jejuni*

Next, we assessed the binding of *C. jejuni* by primary APCs in the PBMC and the effect of 1B5, 1C1 and 2C2. Both Axl^+^ DCs and DC3 showed binding of all three strains of *C. jejuni* tested (Fig. 5C). Interestingly, Axl^+^ DC binding could be blocked by 7-239, but DC3 binding appeared not to be Siglec-1 dependent. In line with our previous data, 2C2 significantly blocked bacterial binding by Axl^+^ DCs, while clones 1B5 and 1C1 did not interfere with *C. jejuni* binding by Axl^+^ DCs (Fig. 5D). No effect of VHH was observed on the binding by DC3. Moderate levels of binding were observed by classical (CD14^+^, CD16^-^) and intermediate monocytes (CD14^+^, CD16^+^) of strains GB31 and GB14 and lower levels of binding of GB23 (Fig. S5B). Similar to DC3, binding by classical and intermediate monocytes was not mediated by Siglec-1 (Fig. S5C, E). In summary, we have shown that our VHH bind to primary DCs expressing Siglec-1 and that VHH 2C2 specifically blocks *C. jejuni* binding to Siglec-1.

### VHH 2C2 blocks Siglec-1 mediated SARS-CoV-2 binding

Siglec-1 is known to capture viruses such as SARS-CoV-2 via binding to sialylated gangliosides or glycoproteins present in the viral envelope [24, 30]. Here we tested binding of WT recombinant spike protein (S1+S2 trimer) to cells. BWSn cells were pre-incubated with either 500 nM VHH or 100 nM antibody prior to incubation with recombinant spike protein (Fig. 6A). VHH clone 1B5 and 1C1 did not affect spike protein binding whereas VHH 2C2 blocked binding to the same degree as mAb 7-239 (Fig. 6A). To investigate whether the VHH could block Siglec-1-mediated binding of SARS-CoV-2, TSn and moDC cells were pre-incubated with 500 nM of VHH or 100 nM of mAb 7-239 before incubation with wild-type SARS-CoV-2 (hCoV-19/Italy). Viral binding was assessed by the presence of SARS-CoV-2 ORF1b RNA in cell lysates (Fig. 6B-C). While significant amounts of virus binding binding was detected with VHH 1B5 and 1C1, 2C2 completely blocked SARS-Cov-2 binding to TSn cells (Fig. 6B). Similarly, VHH 1B5 marginally affected SARS-Cov-2 binding to moDCs, VHH 2C2 significantly suppressed it, comparable to 7-239 (Fig. 6C). In contrast to TSn cells, moDC also express DC-SIGN a known co-receptor for SARS-CoV-2, which could mediate virus binding when Siglec-1 binding is inhibited (Fig. 6c) [26]. Taken together, we have shown that VHH 2C2 blocks SARS-Cov-2 binding to Siglec-1.

**Fig. 6.**
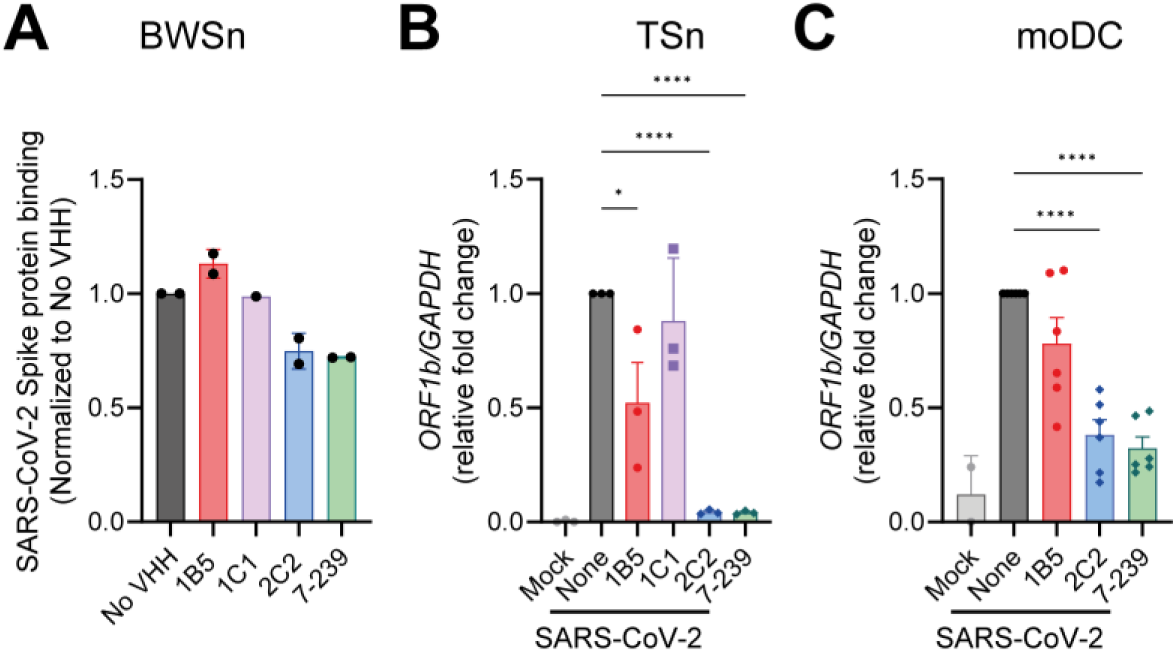
Siglec-1-specific VHH 2C2 blocks SARS-CoV-2 binding. **A)** Recombinant spike protein (S1+S2 trimer) binding to BWSn cells after VHH/antibody pre-incubation as detected by flow cytometry. **B)** qPCR of SARS-CoV-2 ORF1b from cell lysates of Siglec-1 expressing THP-1 cells (TSn) or **C)** moDCs. Cells were pre-incubated with VHH or antibody before incubation with SARS-CoV-2 virus for 4 hours. Data is normalized to GAPDH. B) Representative data is shown for n=3 experiments. C) Data is shown for 6 donors (one way ANOVA: *\*p<0.05, ****p<0.0001*).

## Discussion

To the best of our knowledge, this is the first study to report VHH that allosterically enhance or orthosterically block ligand binding to the sialic acid-binding immunoglobulin-like lectin Siglec-1/CD169. Remarkably, we show that VHH clones 1B5 and 1C1 increase ligand binding, including ganglioside liposomes and *C. jejuni*, to Siglec-1 on moDCs. In contrast, VHH clone 2C2 effectively blocks Siglec-1 ligand interactions and 2C2-mediated inhibition of sialylated pathogen binding such as *C. jejuni* by moDCs, resulted in decreased inflammatory cytokine production by APCs. Furthermore, we reveal for the first time that Axl^+^ DCs can bind *C. jejuni* in a Siglec-1-dependent manner, which was also blocked by 2C2. These results highlight the potential of VHH as effective modulators capable of enhancing or blocking pathogen binding to Siglec-1 on primary cells.

To illuminate how our VHH differentially affect Siglec-1 interactions, we generated AlphaFold3 models of VHH 1B5, 1C1 and 2C2 binding to the first two domains of Siglec-1 and confirmed the results with epitope mapping experiments for 2C2. Although only the terminal Ig-like V-set domain of mouse Siglec-1 has been crystallized, AlphaFold3 could fold the Ig-like V-set domain of human Siglec-1 with a high confidence and superimposition of the structure onto mouse Siglec-1, revealed an overall similar structure [55]. Interestingly, AlphaFold3 docked 2C2 directly onto the ligand binding site in a head-on orientation in which R97 is inserted between CDR1 and CDR3 of 2C2. R97 is crucial for sialic acid binding to Siglec-1, as the arginine forms a salt bridge with the carboxylate of sialic acid and thus, the mutant receptor Siglec-1-R97A no longer binds sialylated ligands [55]. Binding of 2C2 to the Ig-like V-set domain of the receptor was confirmed with recombinant Siglec-1-D1. Moreover, a single point mutation, R97A, in the full length receptor was sufficient to abrogate 2C2 binding, pin-pointing the interaction interface to the ligand binding site and elucidating 2C2’s capacity to block Siglec-1/ligand interactions.

AlphaFold3 was unable to confidently dock 1B5 and 1C1 onto the Ig-like V-set domain of Siglec-1. Introduction of the Ig-like C2-set domain 1 was sufficient to generate models with a high confidence. In the AlphaFold3 model, the Ig-like V-set domain of Siglec-1 is connected to the Ig-like C2-set domain 1 by a short linker. The interdomain interface is further stabilized by a predicted disulfide bridge between Cys17 of V-set domain with Cys147 of the C2-set domain. The AlphaFold3 models suggest that 1B5 and 1C1 target a binding site created by the first two domains of Siglec-1, opposite the ligand binding site. These findings are supported by binding data using recombinant protein and antibody displacement experiments as both VHH required Siglec-1 domain 1 and 2 for binding, but were blocked by 7D2, which exclusively binds to domain 1. Nonetheless, we cannot rule out binding to solely domain 2 as we could not express this domain on its own. Unlike conventional antibodies, VHH have been reported to engage their target antigens in a diverse set of conformations [56]. In the AlphaFold3 model, 1B5 assumed a side-on orientation in its interaction with Siglec-1-D1-2 driven by CDR3 and the VHH framework, while 1C1 was bound in a head-on orientation.

The question remains how the binding of 1B5 and 1C1 VHH can enhance ligand binding despite not binding directly to the ligand binding site. VHH can act as conformational stabilizers locking target proteins in distinct conformations such as inactive or active states of GPCRs, or allosterically regulating enzyme activity [46, 59]. In the case of Siglec-7 it has been shown that the CC’ loop in the Ig-like V-set domain has an ‘open’ form in the native state that undergoes a large conformational change, to a ‘closed’ orientation, upon binding of the ligand GT1b [60]. However, for Siglec-1 no large scale conformational changes have been observed upon ligand binding [55, 61]. Alternatively, due to the high levels of sialic acid found on the cell surface Siglecs can bind ligands on the same membrane in so called *cis*-interactions, which need to be unmasked for the receptor to engage in *trans*-interactions with ligands on opposing cells [10]. Siglec-1 is the largest member of the CD33-related Siglec family with 17 extracellular Ig domains, which in the literature has been suggested to enable the receptor to escape inhibitory *cis-*interactions and pre-dominantly engage ligands in *trans* [10]. Nonetheless, it has been shown that soluble Siglec-1 ligand binding can be enhanced by de-sialylation of various cell types expressing Siglec-1, thereby unmasking the receptor [15–17]. Here, a significant increase in GM3 liposome and 7-239 antibody binding was observed upon neuraminidase treatment of Siglec-1 expressing BWSn cells, which indicates that Siglec-1 is engaged in *cis*-interactions on these cells. It could be speculated that VHH 1B5 binding to Siglec-1 affects the overall rigidity of the receptor making it less capable of engaging in *cis*-interactions and thereby increasing soluble ligand binding. Neuraminidase treatment did significantly reduce the effect of VHH 1B5, suggesting that *cis*-interactions are involved. The degree of *cis*-interactions may vary between various Siglec-1^+^ cells and may potentially influence the effect of 1B5 on ligand binding. While we observed strong enhancing effects of 1B5 and 1C1 on BWSn and TSn cells, this was not observed with VHH 1B5 binding to Axl^+^ DCs and further studies will be necessary to determine whether Axl^+^ DCs have less *cis-*interacting Siglec-1.

Here, we investigated the effects of 1B5, 1C1 and 2C2 on Siglec-1 ligand binding in the context of bacterial and viral pathogens. Pathogens are known to incorporate host-like molecules to circumvent or manipulate the host immune response in a process called molecular mimicry [62]. In the case of *C. jejuni* infections, molecular mimicry is thought to be the cause of GBS, a deadly post-infectious neuropathy, resulting from *C. jejuni* antibodies cross-reacting with gangliosides on host peripheral nerves [36, 63]. Sialylated LOS structures are known to enhance endocytosis of *C. jejuni* and subsequent translocation across intestinal epithelial cells [64, 65], but also mediate binding to Siglec-1 and uptake by macrophages and mouse DCs, resulting in enhanced cytokine production [23, 39–41]. In line with this, we report *in vitro* binding of GBS-related strains GB14, GB23 and GB31 by TSn and moDCs in a Siglec-1-dependent manner, which could be either enhanced by 1B5/1C1 or blocked by 2C2 resulting in reduced IL-6 levels by moDCs. In a murine infection model, sialylated *C. jejuni* was shown to induce the production of GBS-associated cross-reactive antibodies targeting gangliosides and blocking murine Siglec-1 was sufficient to prevent the production of cross-reactive antibodies in this model [42]. We show for the first time that Siglec-1^+^ Axl^+^ DCs but not other DC subsets bind to GBS-associated *C. jejuni* strains in a Siglec-1-dependent manner and that 2C2 specifically blocks this interaction. While a potential role for Axl^+^ DCs in the development of GBS is not known, further research into the therapeutic application of Siglec-1 blocking is warranted.

Multiple enveloped viruses, including SARS-Cov-2, are known to contain sialylated gangliosides and exploit binding to Siglec-1 to infect cells of the myeloid compartment [26, 27, 66]. SARS-CoV-2 attachment to Siglec-1 not only enables infection of Siglec-1^+^ cells, but also trans-infection of other myeloid and lymphoid cells, contributing to viral dissemination [24, 28, 31]. Instead of using neutralizing antibodies for each specific virus strain, blocking antibodies to Siglec-1 have been developed as a pan-antiviral therapy [67]. Here we show that SARS-Cov-2 binding to TSn and moDCs was efficiently blocked by VHH 2C2. Since VHH are smaller than antibodies and penetrate tissues better, Siglec-1 blocking VHH could be used as an alternative option for therapeutic pan-antiviral applications.

## Conclusion

In this study, we have identified Siglec-1 binding VHH that either allosterically enhance or orthosterically block ligand binding. We show that these VHH modulate Siglec-1 interactions within the context of sialylated liposomes and pathogens (*C. jejuni* and SARS-CoV-2) thereby enabling the further study of Siglec-1’s role within infectious diseases. Finally, the identified Siglec-1-specific VHH are interesting modalities for further investigation as potential therapeutics, as VHH can easily be engineerd to generate multivalent proteins with enhanced affinities, or be made bi-specific to block multiple pathogen receptors.

## Acknowledgements

This work was supported by Dutch Cancer Society grants (2019-2/12802 , 2025-AF-1 / 17146 and 2021-2/14093), PPP grants awarded by Health Holland, Top Sector Life Sciences & Health, to stimulate public-private partnerships (Amsterdam UMC and KWF PPS VU2022-14453) and NWO ENW KIEM (ENPPS.KlEM.019.008) to J.M.M.H., ZonMw Open Competitie (09120232310030) to J.M.M.d.H., and T.B.H.G., Amsterdam institute for lnfection and Immunity Collaboration grant to A.J.A. and T.B.H.G., NWO Veni ZonMw (09150162010163), Health Holland, Top Sector Life Sciences & Health in collaboration with Aidsfonds (LSHM19101/P-44802) and the Amsterdam UMC (2019-1167) to T.B.H.G and N.A.K..Stichting Cancer Center Amsterdam (CCA2022-9-83), and KWF Dutch Cancer Society (16429) to A.J.A. We would like to thank Dr. Peter Steinberger from the Medical University of Vienna for providing the BW5147 overexpressing Siglec-1 and dr. Johannes Stöckl for providing the 7-239 hybridoma. We acknowledge the Microscopy and Cytometry Core Facility (MCCF) at the Amsterdam UMC for providing assistance with confocal microscopy and flow cytometry experiments. This work benefited from access to the Netherlands Cancer Institute (NKI) protein production facility, an Instruct-ERIC centre. Financial support was provided by Instruct-ERIC (PID36153, PID24108).

## Data Availability

All study data are included in the article and supporting information.

## Author contributions

H.J.B., A.J.A.,S.M., N.A.K., T.B.H.G. and J.M.M.H. designed the study and experimental approaches. H.J.B., A.J.A.,S.M., N.C.C.T.D., E.A.G, R.G.B., V.A.L.K., J.G.C.S., J.L.H., K.O., D.A.M.H. and A.F. carried out the experiments. H.J.B., A.J.A, N.C.C.T.D., R.G.B., N.S., A.M., P.H.N.C, G.D., M.J.S., R.H. and E.M. generated and provided reagents. H.J.B., A.J.A, S.M., N.A.K., T.B.H.G., Y.V.K. and J.M.M.H. supervised the research. H.J.B. and A.J.A. prepared the figures. H.J.B., A.J.A and J.M.M.H. wrote the manuscript with input from all co-authors. All authors read and approved the final manuscript.

## Patent

WO2024096735

