## Supplemental figures for "Identification of allo- or orthosteric VHH/single-domain antibodies that enhance or block pathogen binding to Siglec-1 on dendritic cells"

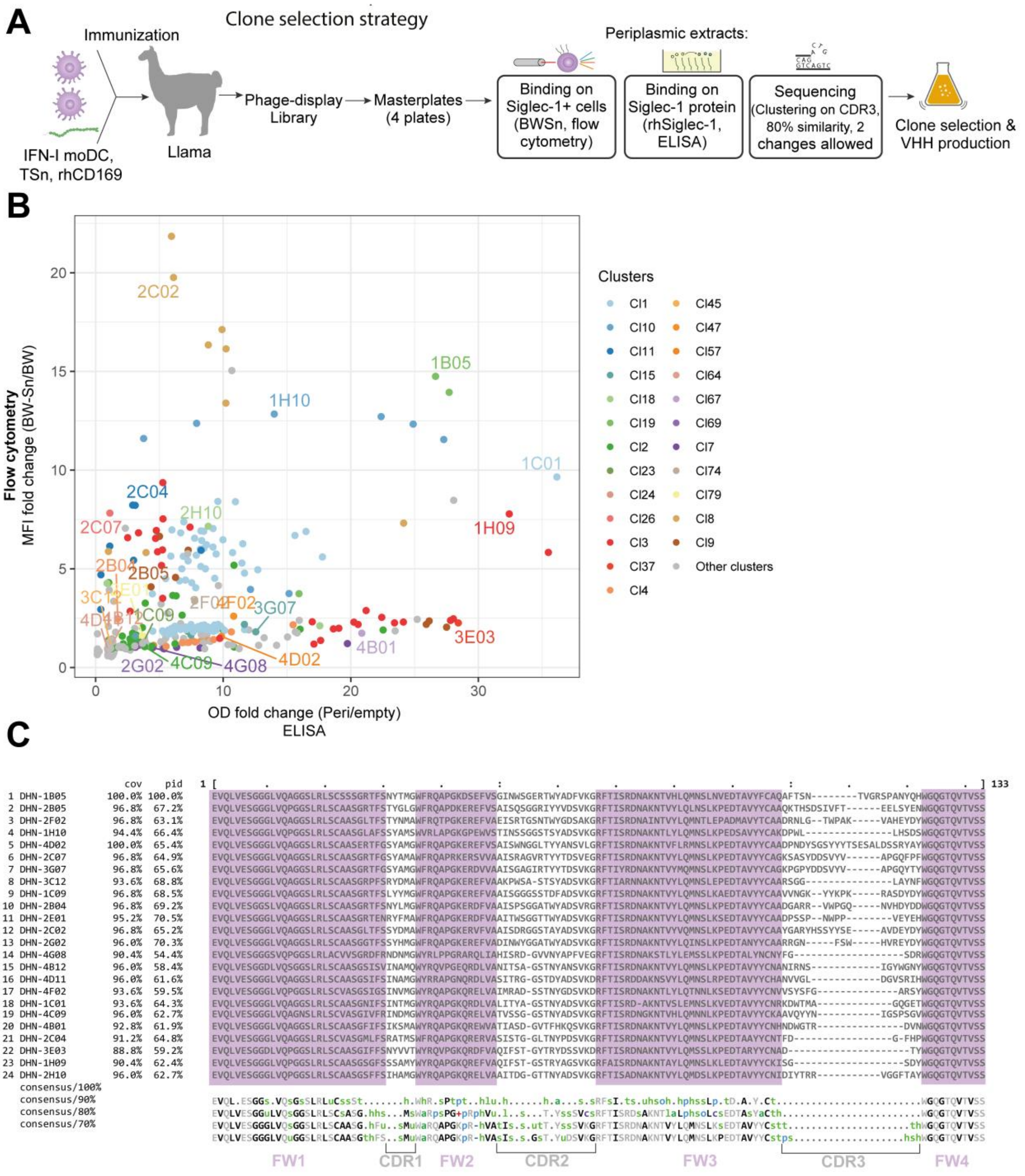


**Figure S1 Selection of Siglec-1-binding VHHs. A)** Schematic overview of VHH immunization and clone selection strategy. **B)** VHH periplasmic binding data to (y-axis) BWSn cells, as determined by flow cytometry, or (x-axis) binding to recombinant protein, as determined by ELISA. Selected clones are annotated. **C)** Sequence alignment of selected VHH clones. Framework regions are highlighted purple. Consensus sequences are shown below.

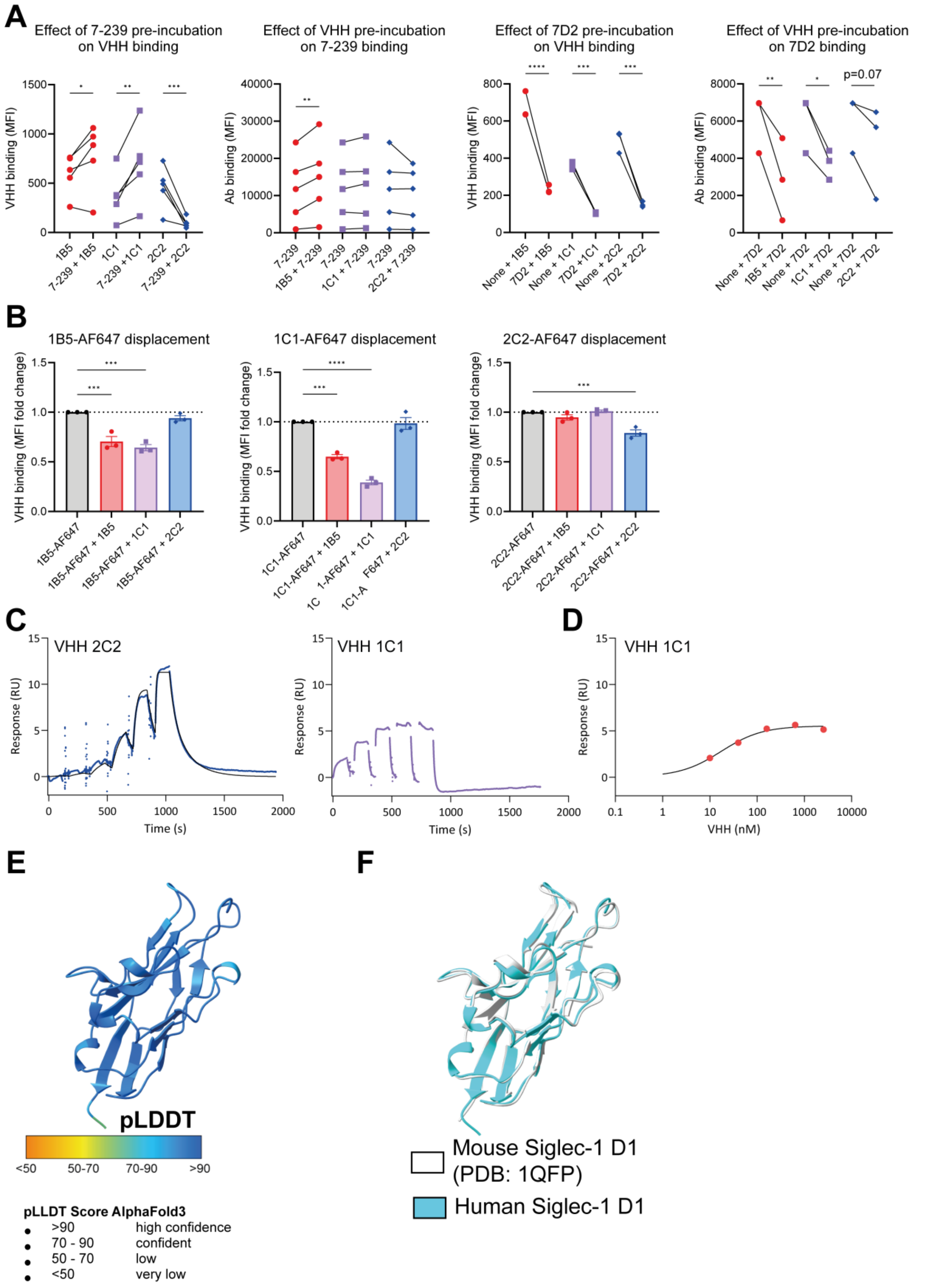
A
Effect of 7-239 pre-incubation on VHH binding
Effect of VHH pre-incubation on 7-239 binding
Effect of 7D2 pre-incubation on VHH binding
Effect of VHH pre-incubation on 7D2 binding
VHH binding (MFI)
Ab binding (MFI)
p=0.07
B
1B5-AF647 displacement
1C1-AF647 displacement
2C2-AF647 displacement
VHH binding (MFI fold change)
C
VHH 2C2
VHH 1C1
Response (RU)
Time (s)
D
VHH 1C1
VHH (nM)
E
pLDDT
<50
50-70
70-90
>90
pLLDT Score AlphaFold3
>90 high confidence
70 - 90 confident
50 - 70 low
<50 very low
F
Mouse Siglec-1 D1 (PDB: 1QFP)
Human Siglec-1 D1

G

| VHH/Siglec-1: | 2c2/Siglec-1D1 | 1B5/Siglec-1 D1-2 | 1C1/Siglec-1 D1-2 |
|---|---|---|---|
| pTM | 0.80 | 0.86 | 0.80 |
| iPTM | 0.86 | 0.88 | 0.86 |

H

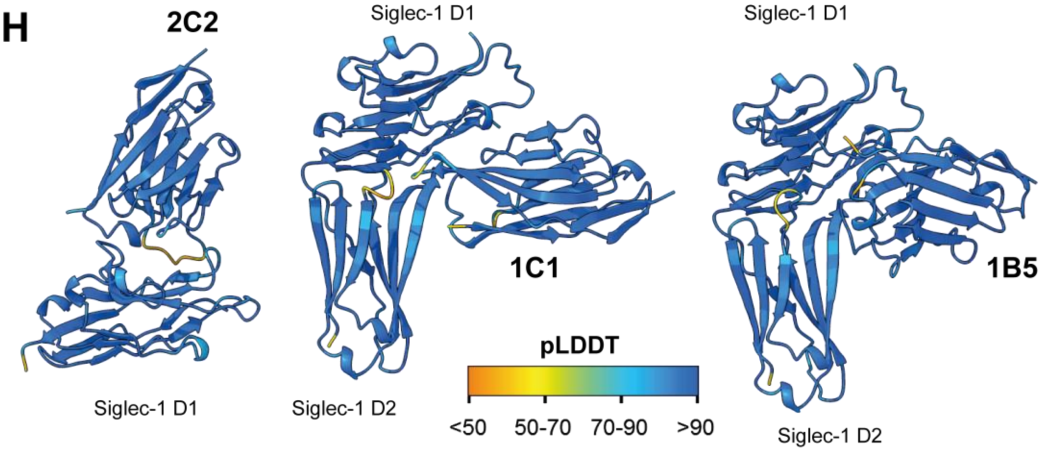


**Figure S2 Epitope mapping of Siglec-1-binding VHH.** Effect of pre-incubation of BWSn cells with antibody **A)** 7-239 or 7D2 on VHH binding or the reverse by flow cytometry. Data is shown as mean fluorescent intensity. **B)** Binding data of AF647 labelled VHH after competition binding with non-labelled VHH to BWSn cells. **C)** Binding of VHH 2C2 and 1C1 to immobilized recombinant Siglec-1 (Ser20-Gln-1641) as measured by SPR. Data was fitted using a 1:1 kinetic binding model fit for 2C2 or for 1C1 **D)** show in an R-max plot and fitted for equilibrium binding. **E)** Alphafold3 fold of human Siglec-1 domain 1. Model is colored according to the predicted local distance difference test scores (pLDDT) **F)** Alphalfold3 fold of human Siglec-1 superimposed on the crystal structure of mouse Siglec-1 domain1 (PDB: 1QFP). **G)** Table of predicted template modeling (pTM) and interface predicted modeling (ipTM) scores per fold shown in show in H. **H)** Alphafold3 model of VHH clone 2C2 docked to Siglec-1 domain 1 or VHH clones 1B5 and 1C1 docked to Siglec-1 domain 1-2. Models are colored according to the pLDDT scores. Data in A and B is shown as mean ± SEM of (t-test: $^{*}p<0.05$, $^{**}p<0.01$, $^{***}p<0.001$, $^{****}p<0.0001$, one way ANOVA: $^{***}p<0.001$, $^{****}p<0.0001$).

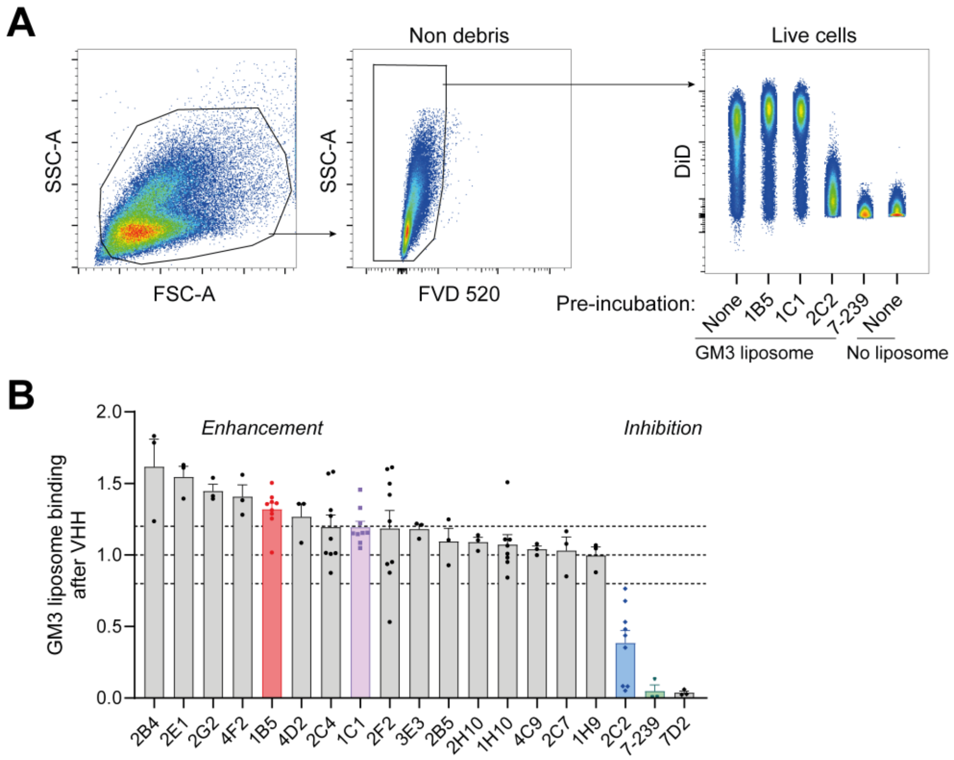


**Figure S3 VHH mediated modulation of ganglioside liposome binding to BWSn. A)** Gating strategy to identify DiD+ cells after pre-incubation of Siglec-1 expressing BWSn with VHH or antibody followed by incubation with GM3 liposomes. **B)** GM3 liposome binding after VHH or antibody pre-incubation as determined by flow cytometry. Data is normalized to a no VHH control and shown as mean ± SEM of at least n=3 experiments.

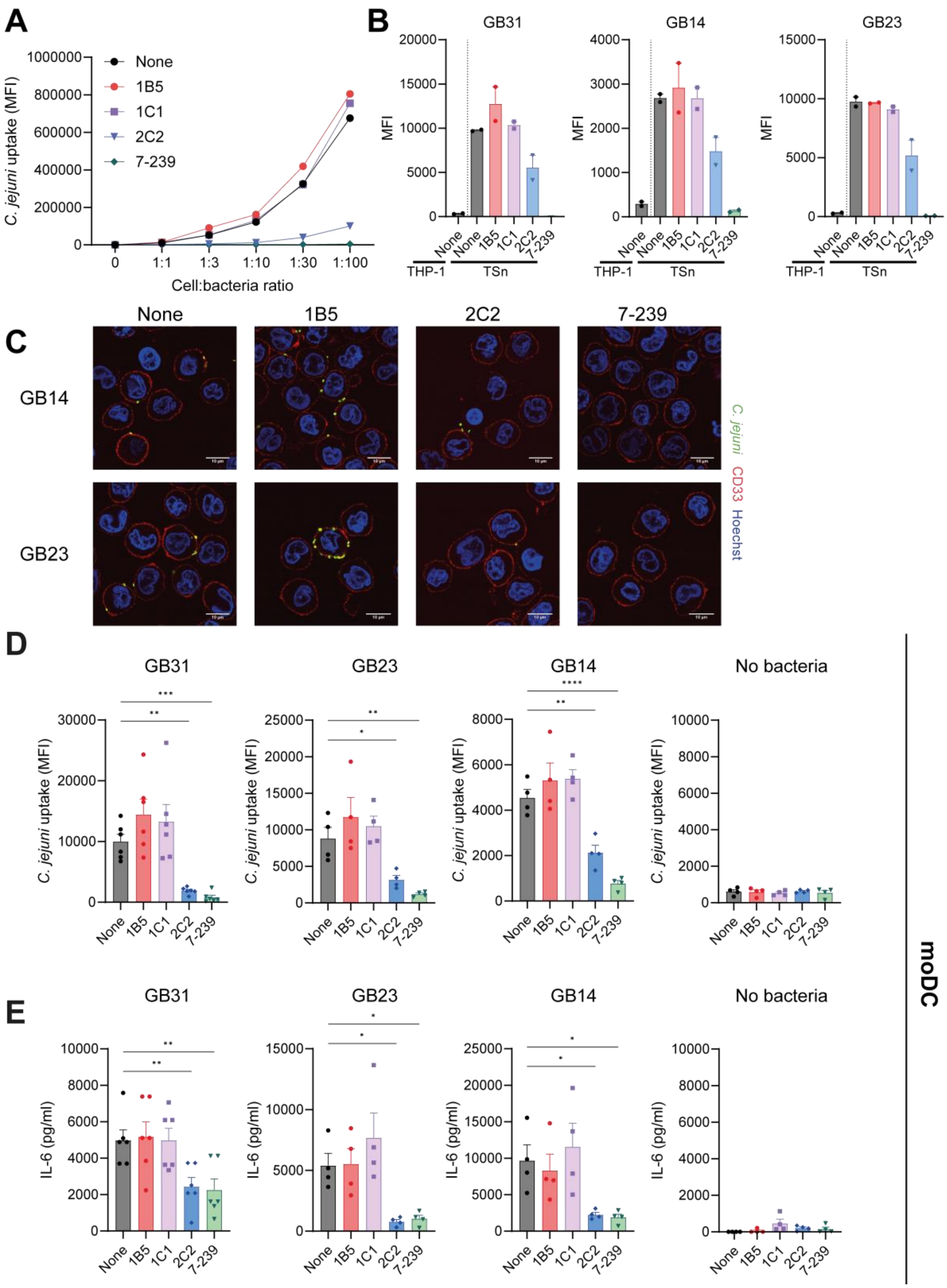


**Figure S4 VHH modulate uptake of *C. jejuni*. A)** *C.jejuni* strain GB31 uptake by TSn at increasing cell to bacteria ratios after pre-incubation with VHH or antibody. **B)** Uptake of *C. jejuni* strains GB31, GB23 and GB14 by THP-1 and TSn cells pre-incubated with VHH or antibody as determined by flow cytometry. Data is as mean ± SD of two experiments. **C)** Fluorescent microscopy images of FITC-labelled GB14 and GB23 *C. jejuni* binding to TSn cells pre-incubated with 7-239 or VHH. TSn membranes detected with anti-CD33-AF647 antibody. Nucleus is stained with Hoechst. **D)** Uptake of *C. jejuni* strains GB31, GB23 and GB14 by moDCs pre-incubated with VHH or antibody as determined by flow cytometry. Data is shown as mean ± SEM of 4 to 6 donors. (one way ANOVA: ***p<0.01, ***p<0.001, ****p<0.0001*) **E)** IL-6 secretion by moDCs, pre-incubated with VHH or antibody, followed by a 6 hour incubation with *C. jejuni* strains GB31, GB23 and GB14 at 37°C. IL-6 amounts were determined by ELISA. Data is shown as mean ± SEM of 4 to 6 donors (one way ANOVA: **p<0.05, **p<0.01, ****p<0.0001*).

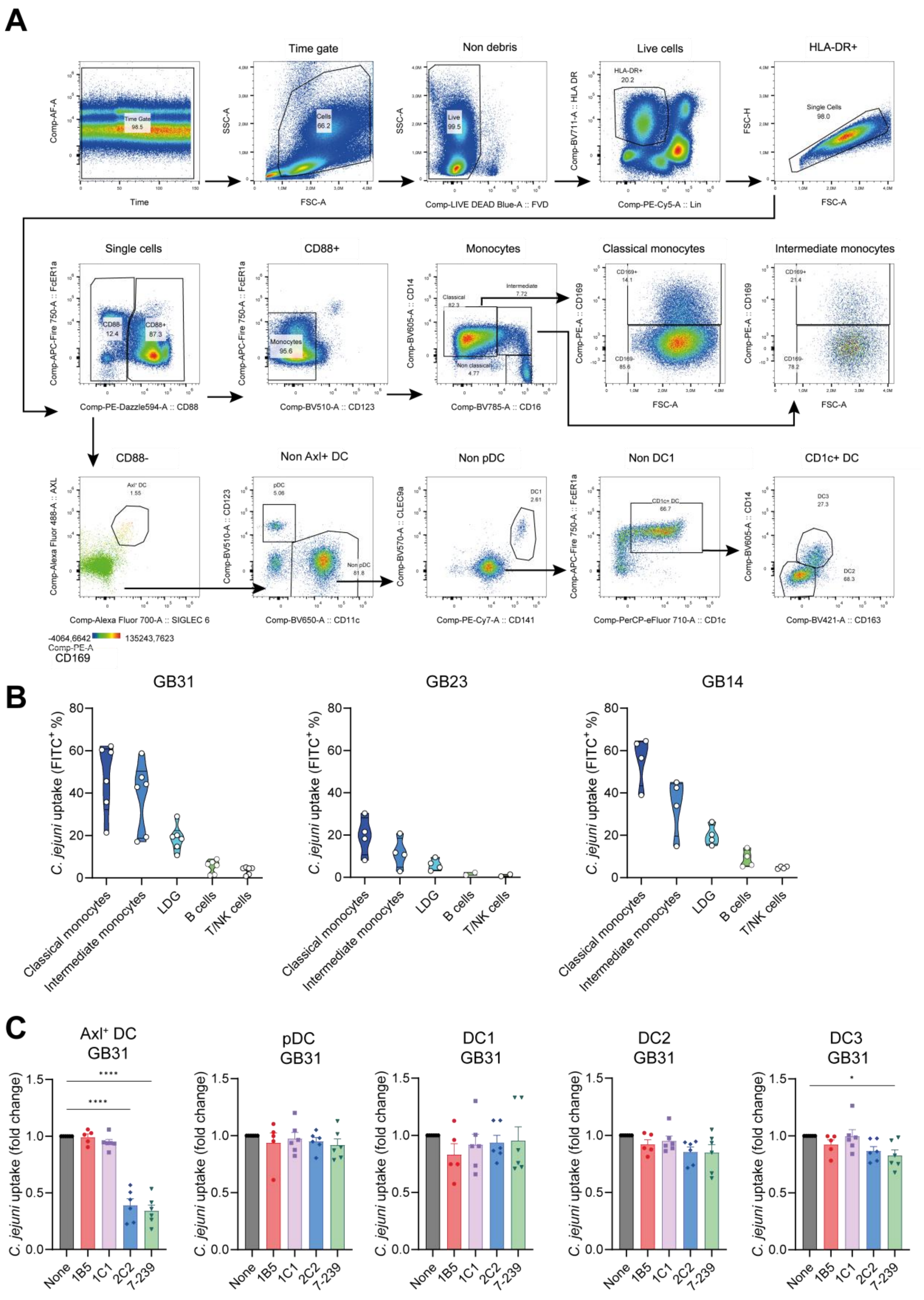

A
Time gate
Non debris
Live cells
HLA-DR+
Comp-AF-A
Time
Time Gate 98.5
SSC-A
FSC-A
Cells 66.2
Live 99.5
Comp-LIVE DEAD Blue-A :: FVD
Comp-BV711-A :: HLA DR
HLA-DR+ 20.2
Comp-PE-Cy5-A :: Lin
FSC-H
Single Cells 98.0
Single cells
CD88+
Monocytes
Classical monocytes
Intermediate monocytes
Comp-APC-Fire 750-A :: FcER1a
Comp-PE-Dazzle594-A :: CD88
CD88- 12.4
CD88+ 87.3
Monocytes 95.6
Comp-BV510-A :: CD123
Comp-BV605-A :: CD14
Comp-BV785-A :: CD16
Classical 82.3
Intermediate 7.72
Non classical 4.77
Comp-PE-A :: CD169
CD169+ 14.1
CD169- 85.6
CD169+ 21.4
CD169- 78.2
CD88-
Non Axl+ DC
Non pDC
Non DC1
CD1c+ DC
Comp-Alexa Fluor 488-A :: AXL
Axl+ DC 1.55
Comp-Alexa Fluor 700-A :: SIGLEC 6
Comp-BV510-A :: CD123
pDC 5.06
Non pDC 81.8
Comp-BV650-A :: CD11c
Comp-BV570-A :: CLEC9a
DC1 2.61
Comp-PE-Cy7-A :: CD141
Comp-APC-Fire 750-A :: FcER1a
CD1c+ DC 66.7
Comp-PerCP-eFluor 710-A :: CD1c
Comp-BV605-A :: CD14
DC3 27.3
DC2 68.3
Comp-BV421-A :: CD163
-4064,6642
135243,7623
Comp-PE-A
CD169
B
GB31
GB23
GB14
C. jejuni uptake (FITC+ %)
0
20
40
60
80
Classical monocytes
Intermediate monocytes
LDG
B cells
T/NK cells
C
Axl+ DC GB31
pDC GB31
DC1 GB31
DC2 GB31
DC3 GB31
C. jejuni uptake (fold change)
0.0
0.5
1.0
1.5
****
****
*
None
1B5
1C1
2C2
7-239

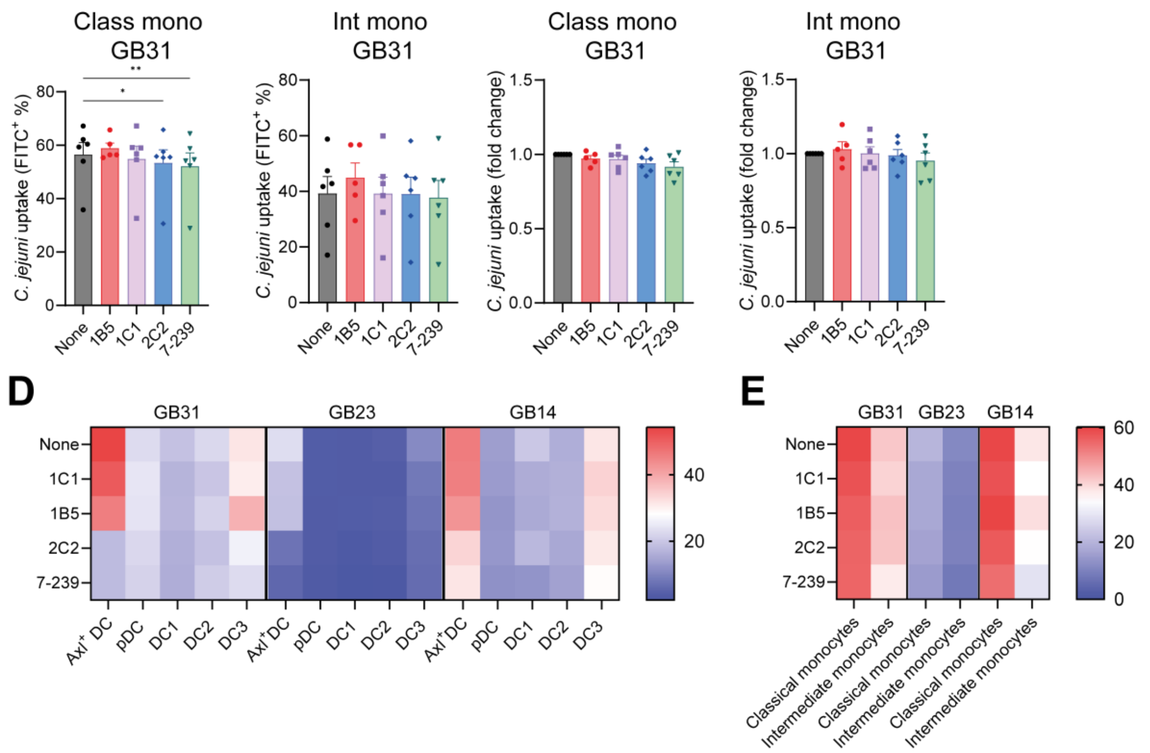


**Figure S5 VHH 2C2 blocks uptake of *C. jejuni* by primary cells. A)** Conventional gating identifies classical monocytes, intermediate monocytes, plasmacytoid DCs (pDC), conventional DC1 (cDC1), conventional DC2 (cDC2), DC3, and Axl+ Siglec-6+ DCs (Axl+ DC). **B)** Percentage FITC+ cells after incubation with *C. jejuni* strains GB31, 23 and 14. Data is shown for 2 to 6 donors. **C)** Uptake of *C. jejuni* strains GB31, GB23 and GB14 by primary cells pre-incubated with VHH or antibody as determined by flow cytometry. Data is normalized to a no VHH control and shown as mean ± SEM of 4 donors. (one way ANOVA: **p=0.05*, ***p<0.01*, *****p<0.0001*). **D)** Heat map of *C. jejuni* strains GB31, 23 and 14 uptake by Axl+ DC, pDC and DC1-3 cells or **E)** classical or intermediate monocytes after pre-incubation with VHH or antibody shown as % of population positive for indicated GB strain.
